# Biofilm self-patterning: mechanical forces drive a reorientation cascade

**DOI:** 10.1101/2021.05.11.440221

**Authors:** Japinder Nijjer, Changhao Li, Qiuting Zhang, Haoran Lu, Sulin Zhang, Jing Yan

**Author notes:** Correspondence and requests for materials should be addressed to either or.

## Abstract

In growing active matter systems, a large collection of engineered or living autonomous units metabolize free energy and create order at different length scales as they proliferate and migrate collectively. One such example is bacterial biofilms, which are surface-attached aggregates of bacterial cells embedded in an extracellular matrix. However, how bacterial growth coordinates with cell-surface interactions to create distinctive, long-range order in biofilms remains elusive. Here we report a collective cell reorientation cascade in growing *Vibrio cholerae* biofilms, leading to a differentially ordered, spatiotemporally coupled core-rim structure reminiscent of a blooming aster. Cell verticalization in the core generates differential growth that drives radial alignment of the cells in the rim, while the radially aligned rim in turn generates compressive stresses that expand the verticalized core. Such self-patterning disappears in adhesion-less mutants but can be restored through opto-manipulation of growth. Agent-based simulations and two-phase active nematic modeling reveal the strong interdependence of the driving forces for the differential ordering. Our findings provide insight into the collective cell patterning in bacterial communities and engineering of phenotypes and functions of living active matter.

---

The spatiotemporal patterning of cells is a fundamental morphogenetic process that has profound effects on the phenotypes and functions of multicellular organisms^1–3^. In the prokaryotic domain, bacteria are often observed to form organized multicellular communities surrounded by extracellular matrices^4,5^, known as biofilms^6,7^, which are detrimental due to persistent infections, clogging of flows, and surface fouling, but can be beneficial in the context of wastewater treatment^8^ and microbial fuel cells^9^. During development, biofilms exhibit macroscopic morphological features ranging from wrinkles, blisters, to folds^10–12^. At the cellular scale, recent progress in single-cell imaging has revealed the reproducible three-dimensional architecture and developmental dynamics of biofilms^13–16^. However, how the cellular ordering emerges from individual bacterium trajectories remains poorly understood. In particular, it remains unclear how cell proliferation is coordinated with intercellular interactions in a growing biofilm to elicit robust self-patterning against bacteria’s inherent tendency to grow in an unstructured manner^17–19^. An understanding of how individual cell growth links to collective patterning as a result of self-generated forces can provide insights into the developmental program of biofilms^6^, their physical properties^20^, and the engineering of living and nonliving active-matter analogs^21,22^.

To bridge the gap between interactions at the cellular scale and patterns at the community scale, here we combine single-cell imaging and agent-based simulations to reveal the underlying mechanism for self-patterning in biofilm formed by *Vibrio cholerae*, the causal agent of the pandemic cholera. We observe that biofilm-dwelling bacteria self-organize into an aster pattern, which emerges from a robust reorientation cascade, involving cell verticalization in the core and radial alignment in the growing rim. We reveal that the verticalized core generates directional flow that drives radial alignment of the cells in the periphery, while the radially aligned rim generates compressive stresses that expand the verticalized core, leading to a robust, inter-dependent differential orientational ordering. Based on these findings, we derive a two-phase active nematic model for biofilm self-patterning, which is potentially generalizable to other developmental systems with growth-induced flows^23,24^. Our findings suggest that the self-generated cellular force landscape, rather than chemical signaling or morphogen gradients as often seen in eukaryotic cells^25^, controls pattern formation in biofilms.

## *V. cholerae* biofilms self-organize into aster patterns

We imaged the growth of *V. cholerae* biofilms confined between glass and an agarose gel at single-cell resolution (Fig. 1a). We used a constitutive biofilm producer locked in a high c-di-GMP state^26^ and focused on the biophysical aspects of self-organization. To simplify our studies, we focused on a mutant missing the cell-to-cell adhesion protein RbmA - this strain is denoted as WT* - although our analysis is equally applicable to strains with cell-to-cell adhesion (Extended Data Fig. 1). Using confocal microscopy, the 3D architecture of the biofilms was captured over time from single founder cells to mature biofilms consisting of thousands of cells (Fig. 1b; Supplementary Video 1). An adaptive thresholding algorithm was used to segment individual cells in the 3D biofilm (Extended Data Fig. 2; Supplementary Information Section 1) from which the location and direction of each rod-shaped bacterium were identified (Fig. 1c-f). Strikingly, cells in the basal layer of WT* biofilms *reproducibly* self-organized into an aster pattern, consisting of a core with tilted or “verticalized” cells and an outward splaying rim with radially aligned cells (Fig. 1d; Extended Data Fig. 1).

**Fig. 1.**
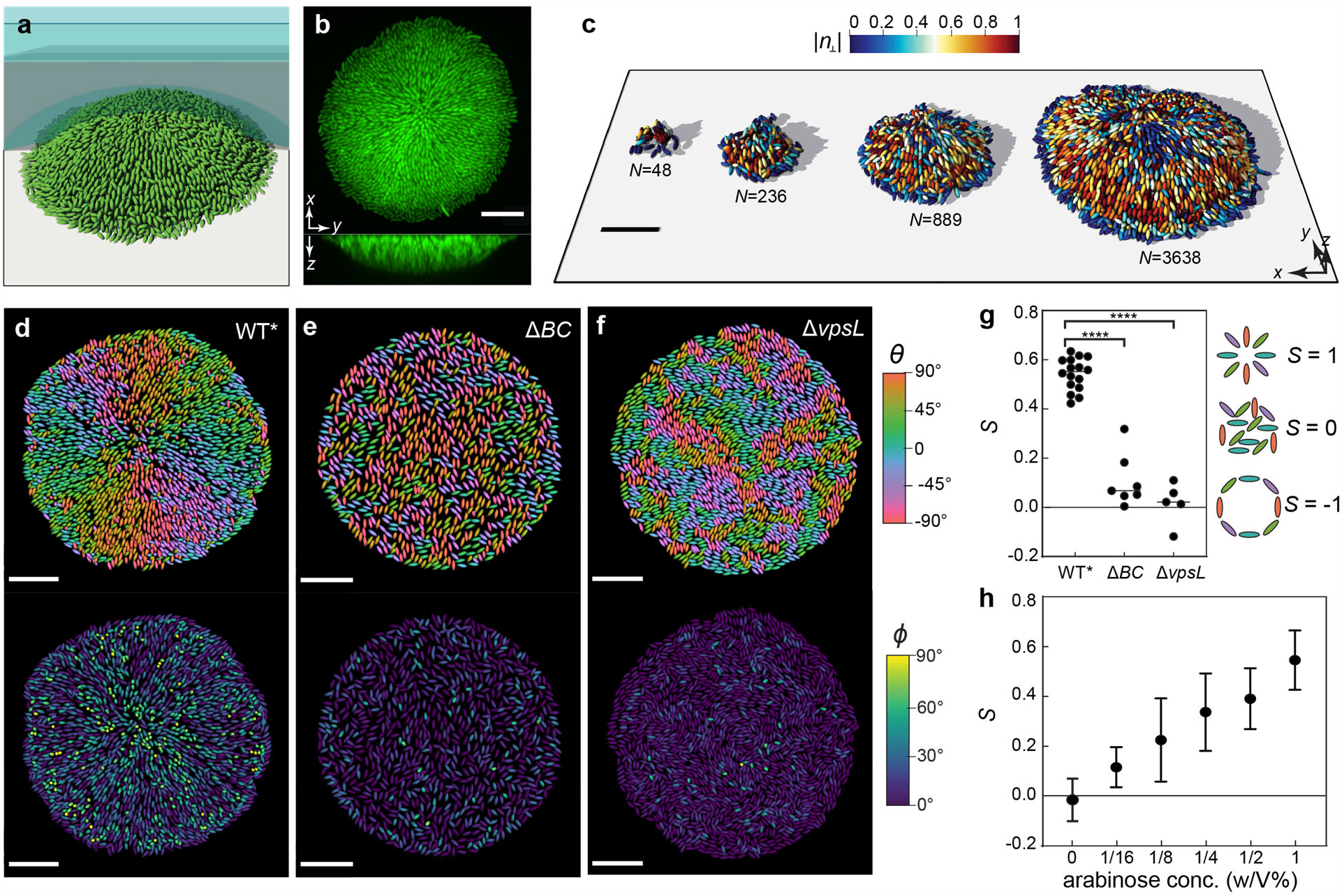
*V. cholerae* biofilms self-organize into aster patterns. **a**, Schematic of the experimental setup, where *V. cholerae* biofilms (green) were grown on a glass surface covered by a hydrogel (blue shaded). **b**, Representative cross-sectional views of a WT* biofilm expressing mNeonGreen. **c**, Single-cell 3D reconstruction of biofilm structures over time with different numbers of cells *N*. **d-f**, Cell orientation color-coded according to each cell’s angle in the basal plane *θ* (*Top*) or the angle it makes with the substrate *ϕ* (*Bottom*), in a biofilm that produces both exopolysaccharides and surface adhesion proteins (WT*; **d**), in a biofilm that only produces exopolysaccharides (Δ*BC;* **e**), and in a bacterial colony with neither exopolysaccharides nor surface adhesion (Δ*vpsL;* **f**). Scale bars, 10 µm. **g**, Radial order parameter *S* quantifying the degree to which cells conform to an aster pattern in the three strains. Data was subjected to ANOVA for comparison of means. ****denotes P<0.0001. **h**, *S* in biofilms in which the expression of *rbmC* is controlled by an arabinose inducible promotor. Error bars correspond to standard deviation.

We quantified the degree of cell ordering in the basal layer using a radial order parameter^27^, 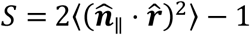, where 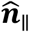 is the projection of the cell direction on the basal plane and 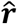 is the unit vector along the radial direction (Fig. 1g). *S* equals 1 for cells that are aligned in an aster, –1 for cells that are aligned in a vortex, and 0 for cells that are randomly oriented. We found that cells in WT* biofilms exhibited a reproducible tendency to align radially (*S* = 0.54 ± 0.07) in the rim. Since previous work has shown that cell-to-surface adhesion controls overall biofilm morphology^12,14,28^, we hypothesized that cell-to-surface adhesion mediates the dynamic core-rim patterning of the biofilm. To test this hypothesis, we deleted the genes encoding cell-to-surface adhesion proteins Bap1 and RbmC^29–31^ (Δ*BC*) and found that the radial order was destroyed in the resulting biofilms and cells assumed random orientations in the basal plane with *S* = 0.11 ± 0.11 (Fig. 1e, g). Concomitant with the disorder was the absence of a verticalized core; most cells in the basal layer were parallel to the substrate. We further confirmed the important role of cell-to-surface adhesion by titrating *rbmC* expression: increasing cell-to-surface adhesion enhanced the self-patterning, resulting in more verticalized cells and stronger radial alignment (Fig. 1h; Extended Data Fig. 3). Furthermore, removing the extracellular matrix by deleting the key *Vibrio* polysaccharide (Δ*vpsL*)^32^ resulted in locally aligned microdomains of horizontal cells without long-range order (*S* = 0.02 ± 0.08; Fig 1f, g), in line with previous studies on growing 2D bacterial colonies^18,19^. These observations suggest that exopolysaccharide production controls a *local* order-to-disorder transition, whereas cell-to-surface adhesion controls a *global* order-to-disorder transition.

To determine the driving forces behind the observed orientational ordering, we extended a previous agent-based model,^33^ taking into account cell-to-cell and cell-to-surface interactions (Supplementary Information Section 2). Our agent-based modeling reproduced the observed aster pattern formation in adherent cells, but not in nonadherent cells, in agreement with experiments (Extended Data Fig. 4; Supplementary Video 2). As the agent-based model only incorporates mechanical interactions, without any biochemical signals, our results suggest that the emergent patterns originate primarily from the mechanical interplay between the cells and between cells and the substrate.

## Surface adhesion drives ordering through differential growth

In molecular liquid crystals, a lower temperature favors order due to the entropic driving force. For out-of-equilibrium systems, such as growing biofilms, the driving force for ordering is more complex. We hypothesized that radial organization arises from the mechanical coupling between cells through their self-generated flow field^34^, inspired by the alignment of rod-shaped objects under fluid shear^35^. Note that biofilm-dwelling cells are nonmotile; flow in this context is generated through cell growth and cell-cell interactions. To test our hypothesis, we tracked cell orientations and trajectories *simultaneously* during biofilm development by using strains expressing a single intracellular punctum (Fig. 2a-e, Extended Data Fig. 5; Supplementary Video 3)^16^. As WT* biofilms grew, cells towards the center tilted away from the substrate, developing a core of verticalized cells that expanded over time (Fig. 2c). The resulting growth-induced flow field had a zero-velocity core (Fig. 2a, d), corresponding to the verticalized cells that project their offspring into the third dimension (Fig. 1d). In contrast, in the nonadherent mutant, the velocity field simply scaled linearly with the radial position. From the measured velocity field, we extracted the *apparent* in-plane proliferation rate *g* (Fig. 2b, Fig. 2d inset). We found that *g* was uniform in the nonadherent biofilm: all cells in the basal layer were predominantly parallel to the substrate and therefore contributed to the basal layer expansion. In contrast, in the WT* biofilm, a growth void (*g* ≈ 0) emerged in the center, with nearly uniform growth in the outer growing rim. Concomitant with the initiation of differential growth, cells aligned in an aster pattern, marked by a growing *S*(*r*) with a rising peak near the edge of the verticalized core (Fig. 2e).

**Fig. 2.**
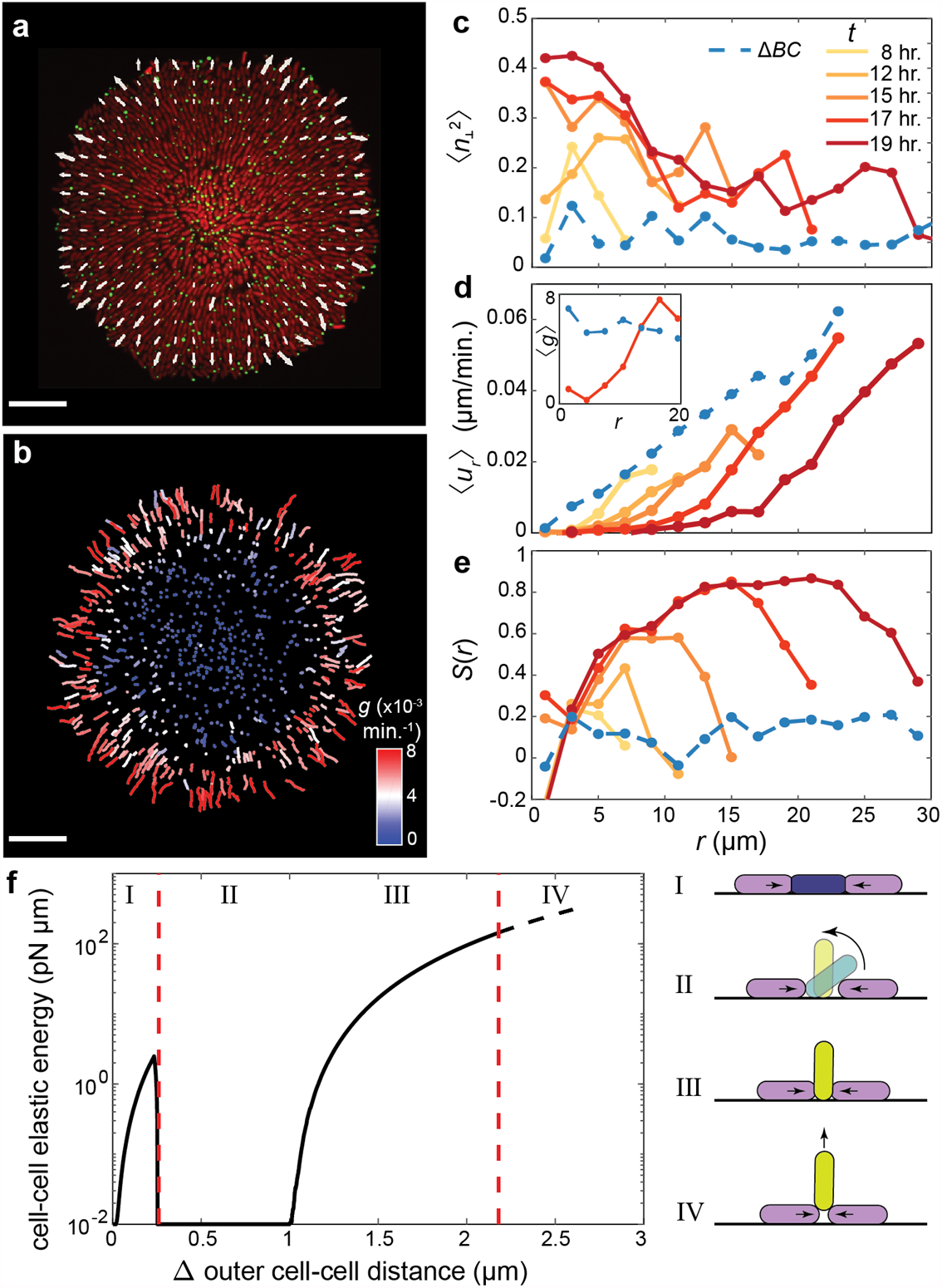
Growth-induced cellular flow and surface anchoring jointly lead to aster formation in biofilms. **a**, Raw image of the basal layer of a biofilm consisting of cells constitutively expressing mScarlet-I cytosolically and mNeonGreen-labelled puncta. Overlain is the velocity field measured from puncta trajectories. **b**, Puncta trajectories colored by the apparent in-plane growth rate *g*. The apparent in-plane growth rate is calculated as *g*(*r*) = (*δ*_*r*_*ru*_*r*_)/*r* in a neighborhood around each cell. Scale bars, 10 µm. **c-e**, Azimuthally averaged degree of verticalization 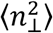 (**c**), radial velocity ⟨*u*_*r*_⟩ (Inset: apparent in-plane growth rate ⟨*g*⟩ × 10^−3^min.^-1^) (**d**), and radial order parameter *S* (**e**), as a function of distance *r* from the center, in the basal layer. The dashed blue lines denote results from the nonadherent mutant. **f**, Results of a reduced problem showing the strain energy due to cell-to-cell contacts in a cell as it is squeezed by two neighbors (black line). The dashed red lines denote the results from stability analyses (Supplementary Information Section 3). Upon increasing compression, the central cell evolves through four phases, which are given schematically.

## A reorientation cascade governs biofilm self-patterning

We hypothesize that a mechanical synergy between cell verticalization, growth-induced flow, and aster pattern formation propels a reorientation cascade for biofilm self-patterning. On one hand, cell-to-surface adhesion coupled with growth-induced mechanical stresses leads to *stably* anchored, verticalized cells in the biofilm center, which results in differentially oriented proliferation. One the other, differential proliferation drives cellular flows that radially align the cells in the rim, which in turn facilitates cell verticalization and core expansion. Below, we analyze the dynamic interplay of these two reorientation processes.

### Step 1

To illustrate the formation and stabilization of the verticalized core, we consider a reduced problem consisting of a spherocylindrical cell that is parallel and adhered to a substrate and squeezed by two neighbors (Supplementary Information Section 3). The resulting energy landscape displays two distinct mechanical instabilities (Fig. 2f). The first instability corresponds to the verticalization event reported earlier^33,36–38^. Briefly, cells in a growing population mechanically push each other, generating pressure. This pressure accumulates and eventually exceeds a threshold, causing cells to rotate away from the substrate (verticalize). The second instability corresponds to the “pinch-off” of these verticalized cells. In this case, neighboring cells generate forces in the out-of-plane direction, causing ejection of the verticalized cells from the substrate. For WT* cells, our analysis shows that pinching a vertical cell off the surface is energetically much more costly than verticalizing a horizontal cell. Therefore, pinch-off is kinetically hindered and verticalized cells can *stably* inhabit the basal layer. The smaller the cell-to-surface adhesion, the smaller the energy difference between the two instabilities (Extended Data Fig. 6) and therefore, the less stable the verticalized cells. The energy difference vanishes in nonadherent cells, resulting in spontaneous ejection of mutant cells upon verticalization. This explains the absence of verticalized cells in the mutant biofilms and bacterial colonies (Fig. 1e, f). In the WT* biofilms, verticalization preferentially occurs near the center where pressure is relatively high, leading to an expanding verticalized core^33^. Since rod-shaped cells grow and divide along their long axes, this spatial segregation of cell orientation leads to spatially patterned differential growth.

### Step 2

Next, we employ active nematic theory^34,39,40^ to elucidate how differential growth can induce radial alignment. Defining the nematic order parameter 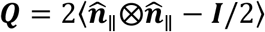 as the head-tail symmetric tensor of cell orientation, mesoscopically averaged over a small region, its evolution in a surrounding flow ***u*** is given by^41^

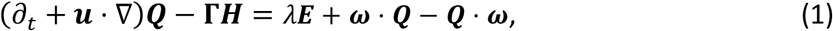

where the right-hand side quantifies the driving force for the rod-shaped particles to rotate within a velocity gradient field. Here 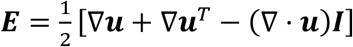 is the traceless strain-rate tensor, 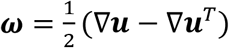 is the vorticity tensor, and *λ* is the flow-alignment parameter. For rod-shaped objects *λ* > 0, corresponding to a tendency for the rods to align with flow streamlines^17^. Finally, the nematic alignment term Γ***H*** relaxes ***Q*** towards a bulk state with minimal angular variation, however, its contribution in biofilms is expected to be negligible since cells are buffered from each other by soft exopolysaccharides (Supplementary Information Section 4). Assuming axisymmetry, the evolution of the cell orientation field is given by^19,34^

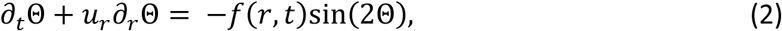

where Θ is the angle between the local orientation field and the radial direction, *f* = (*λr*/4*q*)*∂*_*r*_(*u*_*r*_/*r*) quantifies the aligning torque due to gradients in the flow field, and *q* quantifies the degree of local ordering (Supplementary Information Section 4). From *∂*_t_Θ *∼* −*f* sin(2Θ), we find that a nonzero *f* causes cells to rotate, and the direction of rotation is critically dependent on the sign of *f*.

Unlike passive liquid crystals, biofilm-dwelling cells generate their own velocity field through growth. Assuming uniform density, mass conservation requires ∇ · ***u*** = *g*(*r*). In nonadherent mutant biofilms and bacterial colonies, growth is exclusively in-plane with a uniform growth rate *γ*, resulting in a linear velocity field, *u*_*r*_ = *γr*/2, and thus a vanishing driving force for cell alignment (*f* = 0). Under this condition, cells are simply advected outwards without any tendency to align, leading to a disordered pattern. In contrast, in WT* biofilms, verticalization stabilizes an expanding in-plane growth void, *r*_0_(*t*). This corresponds to a differential growth rate *g*(*r*): 0 for *r* ≤ *r*_0_ and *γ* for *r* > *r*_0_. The resulting velocity field is 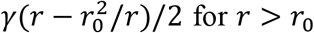, leading to a *strictly positive* driving force for radial alignment, 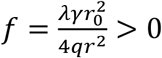, in the outer growing rim. In this case, Θ dynamically approaches 0, characteristic of an aster (Extended Data Fig. 7). In fact, long-range order can be induced whenever a 2D growing bacterial colony deviates from an isotropically expanding pattern, for instance when confined in a rectangular geometry^42,43^ or during inward growth^34^. This model thus reveals that differential growth, established by a verticalized core (*r*_0_ ≠ 0), generates the driving force for radial alignment in a growing biofilm. This driving force vanishes in the absence of a core (*r*_0_ = 0), leading to a disordered phenotype.

## Imposing a growth void reproduces radial ordering

A key prediction of the active nematic theory is that a growth void is *sufficient* to induce radial organization. To test this prediction, we patterned a growth void into an otherwise disordered biofilm. Specifically, we started with a nonadherent biofilm already grown for 17 hours and used a 405 nm laser to selectively kill the cells in the center. The vestiges of the dead cells sustained a growth void (Extended Data Fig. 8), mimicking the verticalized core in the WT* biofilm. Consistent with our model prediction, the proliferating cells aligned radially over time in biofilms with a growth void, whereas biofilms without a growth void remained disordered (Fig. 3a-c). Conversely, our theory predicts that *excess* growth at the biofilm center should lead to *f* < 0 and therefore to vortex formation (Supplementary Information Section 4). Indeed, in another set of experiments, we observed that the nonadherent cells aligned circumferentially when excess growth was introduced at the center (Extended Data Fig. 9). We also quantitatively tested the validity of the model by prescribing a growth void with a fixed size *r*_0_ in a set of simplified 2D agent-based models (Extended Data Fig. 7; Supplementary Video 4). We found that the instantaneous angular velocity of individual cells scaled linearly with sin(2Θ)/*r*^2^ and increasing *r*_0_ led to a quadratic increase in the angular velocity, all in agreement with the theory (Fig. 3d). Note that in both simulations and experiments, the radial order quickly saturated in the patterned biofilm with a fixed *r*_0_, since the aligning force decays with 1/*r*^2^ as cells are advected outward. Thus, a growing *r*_0_(*t*) is necessary to reinforce radial alignment during biofilm expansion. This is indeed the case in WT* biofilms: growth of the outer rim accumulates pressure to generate more verticalized cells and expand the verticalized core, which in turn continuously drives alignment in the outer horizontal cells. To interrogate the mechanical interplay between these reorientation processes, we next develop a minimal physical model coupling verticalization of individual cells to the long-range radial ordering.

**Fig. 3.**
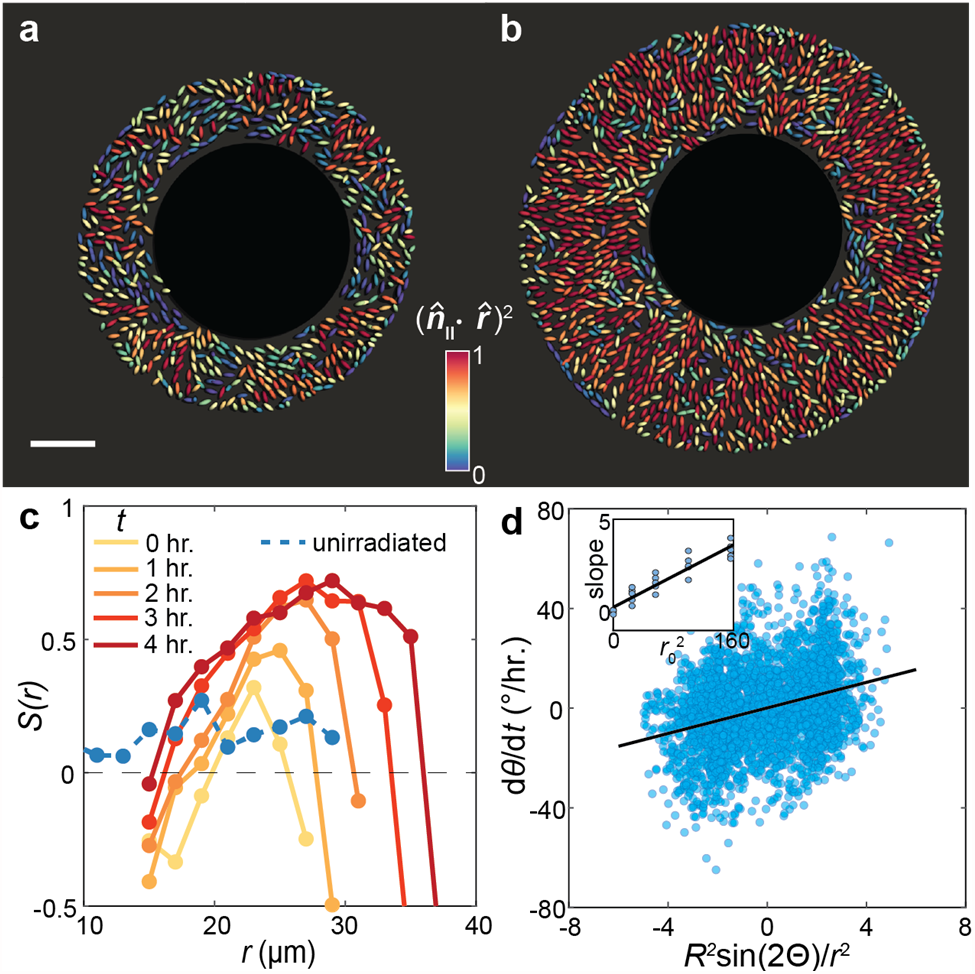
Cell organization can be manipulated by controlling spatial growth patterns. **a, b**, A nonadherent biofilm grown for 17 hours was irradiated using 405 nm laser to induce cell death in a circle of radius 15 μm at the center. Colors denote the degree of radial alignment of individual cells 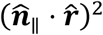 in the irradiated biofilm (colored according to time) and the unirradiated control (blue). **d**, Angular velocity of individual cells from ABSs with a growth void plotted against the predicted nondimensionalized driving force. Inset: Fitted slope from **d** for different growth void sizes *r*_0_ (μm^2^).

## Two-phase model of cell organization

We decompose the biofilm into populations of two phases with vertical and horizontal cells and take the phase fractions to be *ρ* and 1 - *ρ*, respectively. The growth kinetics of the phases are governed by

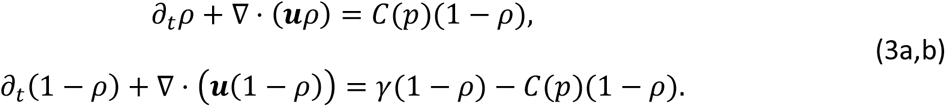

Here we assume that the horizontal-to-vertical conversion is driven by the local pressure *p*, where *C*(*p*) is the conversion rate. We further assume that pressure arises from friction with the substrate ∇*p* = *η****u***, where *η* is the friction coefficient, and that only the horizontal cells generate growth in the basal layer, ∇ · ***u*** = *γ*(1 - *ρ*(***r***)). Combined with Eq. (2), these equations generate a complete continuum description of the dynamics of cell growth and reorientation in biofilms (Supplementary Information Section 4). Numerical solutions of the model quantitatively reproduce the cascade of self-organization events (Fig. 4a-d), showing the intimate spatiotemporal coupling between cell verticalization and radial alignment.

**Fig. 4.**
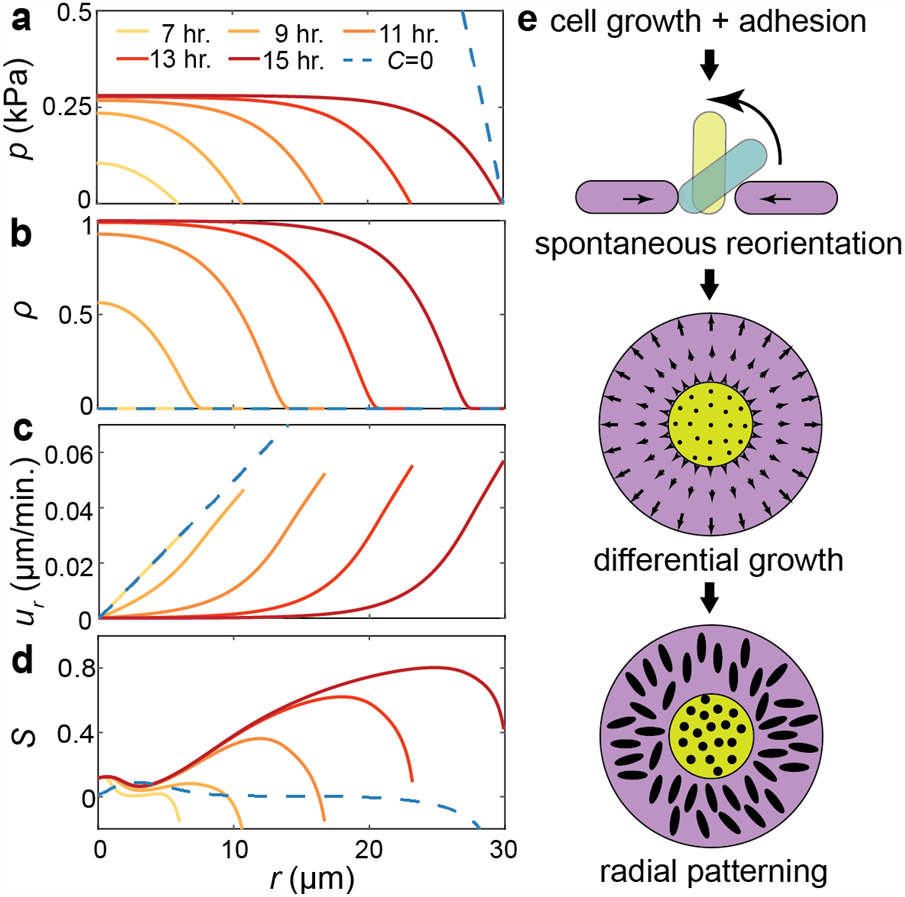
A two-phase active nematic model predicts spontaneous generation of differential proliferation and macroscopic cell organization. **a-d**, Numerical solution of the model consisting of a population of horizontal and vertical cells. The biofilm was initiated with no vertical cells and random in-plane orientations. Evolution of pressure *p* (**a**), fraction of vertical cells *ρ* (**b**), in plane radial velocity *u*_*r*_ (**c**), and radial order parameter *S* (**d**). Curves are colored according to time. Results for a biofilm that cannot sustain verticalized cells (*C* = 0) are shown in blue. **e**, Schematic representation of the biofilm self-patterning process.

Many salient features of the experimental results are recapitulated by the model: for example, *S*(*r*) reaches a maximum near the verticalized core where the driving force is the strongest. Interestingly, the model reveals a frozen core where cells cease to reorganize (compare Fig. 2e and Fig. 4d): as the in-plane velocity goes to zero, the driving force to rotate also vanishes – cells in the core are thus locked as a “fossil record” that memorizes the mechanical history they have experienced. Importantly, the model yields robust results: regardless of the initial conditions and choice of parameters (Extended Data Fig. 10), a WT* biofilm always patterns itself following the sequence shown in Fig. 4e. Our two-phase active nematic model thus elucidates the reproducible mechanical blueprint that guides biofilm development.

## Discussion

To conclude, our results reveal a mechanically driven self-patterning mechanism in bacterial biofilms in which cells synergistically order into an aster pattern. Specifically, we showed that surface adhesion leads to stable cell verticalization, which in turn directs radial cell alignment during surface expansion. Evidently, this inter-dependent differential ordering involves biofilm-wide, bidirectional mechanical signal generation and transmission, in contrast to the biochemical signaling widely observed in other living organisms. In *On Growth and Form*^44^, D’Arcy Thompson wrote: “… growth [is] so complex a phenomena…rates vary, proportions change, and the whole configuration alters accordingly.” Although over a century old, this statement still rings true today. Our two-phase active nematic model provides a mathematical formalism for this statement in the context of bacterial biofilms.

Spontaneous flow generation is a common phenomenon in various developmental systems, including zebrafish embryonic development^24^, ventral furrow formation in *Drosophila*^23^, etc. While flow causes bulk morphological changes in these systems, in biofilms it acts to transmit mechanical forces and drive long-range organization. It is intriguing to contemplate whether the synchronous mechanical coupling between differentially grown cells and the resulting pattern could be generalized to other organisms with anisotropic growth of polarized cells. In a broader context, cell polarity and organization critically underlie collective cell function and normal development, as exemplified by topological defects that mediate 2D-to-3D transitions in motile bacterial colonies^45^ and cell death and extrusion in epithelial layers^46^. Our findings hence shed light on the biomechanical control of cell organization through the spatiotemporal patterning of growth and pave the way to controlling cell organization by encoding synthetic biological circuits or optogenetic manipulation^47^.

## Methods

### Bacterial strains and cell culture

Strains used in this study were derivatives of the *V. cholerae* strain C6706 containing a missense mutation in the *vpvC* gene (*vpvC*^W240R^), which resulted in constitutive biofilm production through the upregulation of c-di-GMP (rugose/Rg strain). For the majority of the results presented in this work, we used a strain in which the gene encoding the cell-to-cell adhesion protein RbmA was deleted to minimize the effects of intercellular adhesion; however, we found that our analysis equally applied to the rugose strain (Extended Data Fig. 1). We primarily worked with two other mutants: 1) Δ*BC* which included additional deletions of *bap1* and *rbmC* genes, and 2) Δ*vpsL* in which a key exopolysaccharide biogenesis gene was deleted in the rugose background (RgΔ*vpsL*). In the absence of *bap1* and *rbmC*, the Δ*BC* mutant cells were unable to adhere to the substrate (referred to as the nonadherent mutant throughout the text). In the absence of *vpsL*, the cells did not properly synthesize exopolysaccharides and consequently, all accessary matrix proteins, which bind to the exopolysaccharide, did not function properly^14^. For velocity field measurements, we used strains containing the μNS protein from the avian reovirus fused to an mNeonGreen fluorescent protein, which were shown to self-assemble into a single intracellular punctum^16,48^. All strains used in the study were also modified to constitutively produce either mNeonGreen or mScarlet-I fluorescent proteins. Mutations were genetically engineered using either the pKAS32 exchange vector^49^ or the MuGENT method^50^. For a full list of strains used, see Table S1. Biofilm growth experiments were performed using M9 minimal media supplemented with 0.5% glucose (w/w), 2 mM MgSO_4_, 100 µM CaCl_2_, and the relevant antibiotics as required (henceforth referred to as M9 media).

Experiments began by first growing *V. cholerae* cells in liquid LB overnight under shaken conditions at 37°C. The overnight culture was back-diluted 30× in M9 media and grown under shaken conditions at 30°C for 2-2.5 hours until it reached an OD_600_ value of 0.1-0.2. The regrown culture was subsequently diluted to an OD_600_ of 0.001 and a 1 µL droplet of the diluted culture was deposited in the center of a glass-bottomed well in a 96-well plate (MatTek). Concurrently, agarose was dissolved in M9 media at a concentration of 1.5-2% (w/V) by microwaving until boiling and then placed in a 50°C water bath to cool without gelation. After cooling, 200 nm far-red fluorescent particles (Invitrogen F8807) were mixed into the molten agarose at a concentration of 1% (V/V) to aid in image registration. Next, 20 µL of the molten agarose was added on top of the droplet of culture and left to cool quickly at room temperature, to gel, and to trap the bacterial cells at the gel-glass interface. Subsequently, 100 µL of M9 media was added on top of the agarose gel, serving as a nutrient reservoir for the growing biofilms. The biofilms were then grown at 30°C and imaged at designated times.

### Image acquisition

Images were acquired using a confocal spinning disk unit (Yokogawa CSU-W1), mounted on a Nikon Eclipse Ti2 microscope body, and captured by a Photometrics Prime BSI CMOS camera. A 100× silicone oil immersion objective (N.A. = 1.35) along with 488 nm, 561 nm and 640 nm lasers were used for imaging. This combination of hardware resulted in an *x*-*y* pixel size of 65 nm and a *z*-step of 130 nm was used. For end-point imaging, biofilms were imaged after 12-24 hours of growth and only the 488 nm channel, corresponding to the mNeonGreen expressing cells, was imaged. For time-lapse imaging, samples were incubated on the microscope stage in a Tokai Hit stage top incubator while the Nikon perfect focus system was used to maintain focus. Images were captured every 30 minutes, and in addition to the 488 nm channel, the 640 nm channel was used to image the fluorescent nanoparticles.

For velocity measurements, cells constitutively expressing mScarlet-I and mNeonGreen-labelled puncta were imaged using a slightly modified procedure. The 488 nm channel, corresponding to the puncta, was imaged every 2-10 minutes while the 561 nm channel, corresponding to the cells, was imaged every 1-2 hours. This procedure allowed us to image the relatively bright puncta with low laser intensity and therefore minimal photobleaching and phototoxicity, as high temporal resolution is required to accurately track puncta motion. To further reduce photobleaching and phototoxicity, we used a *z*-step of 390 nm when imaging the puncta. When imaging the cells, a *z*-step of 130 nm was used in the mScarlet-I channel to sufficiently resolve the 3D position and orientation of the cells. We also restricted our attention to the basal flow field and therefore only imaged the bottom 3 µm of each biofilm. All images shown are raw images rendered by Nikon Elements software unless indicated otherwise.

### Overview of image analysis

Raw images were first deconvolved using Huygens software (SVI) using a measured point spread function. The deconvolved three-dimensional confocal images were then binarized, layer by layer, with a locally adaptive Otsu method. To accurately segment individual bacterium in the densely packed biofilm, we developed an adaptive thresholding algorithm. For more details see Supplementary Information Section 1. Once segmented, we extracted the cell positions by finding the center of mass of each object, and the cell orientations by performing a principal component analysis. The positions and directions of each cell were converted from Cartesian 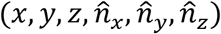 to cylindrical polar 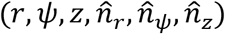 coordinates where the origin was found by taking the center of mass of all of the segmented cells in the (*x, y*) plane. We define the out-of-plane component of the direction vector as 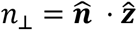 and the in-plane component as 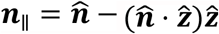, which we normalize as 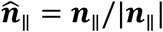. Reconstructed biofilm images were rendered using Paraview.

### Measurement of the growth-induced velocity field

To measure the growth-induced velocity field we used particle tracking velocimetry on the puncta trajectories. The deconvolved puncta images were first registered using Matlab built-in functions. Puncta were then detected by first identifying local intensity maxima in the 3D images, and sub-pixel positional information was found by fitting a parabola to the pixel intensity around the maxima. This procedure was repeated for all frames yielding puncta locations over time which were then connected from frame to frame using a standard particle-tracking algorithm^51^. The radial velocity *u*_*r*_ was calculated by fitting a straight line through the time vs. radial displacement data over a time interval of 1 hr.

### Opto-manipulation of cell growth

Previous work has shown the bactericidal effects of high energy near-UV light^52^; therefore, we used spatially patterned 405 nm light to kill a subset of cells within a biofilm. Specifically, an Opti-Microscan XY galvo-scanning stimulation device with a 405 nm laser was used to selectively illuminate and kill cells within a cylindrical region at the center of the biofilm. We verified cell killing by staining the sample with propidium iodide (Extended Data Fig. 8). The same procedure used to measure the growth-induced velocity field (see above) was applied to the irradiated biofilm and the control to measure cell orientation and trajectory dynamics simultaneously.

### 3D agent-based simulations

Building on the agent-based simulations developed by Beroz *et al*.^33^ and others^37,53,54^, we modelled cells as spherocylinders with a cylinder of length *L*(*t*) and two hemispherical caps of radius *R*. The growth of each cell was assumed to be unidirectional and exponential, where the growth rate *γ* was normally distributed with a mean of *γ*_0_ and a standard deviation of 0.2*γ*_0_. Here noise was added to account for the inherent stochasticity in cell growth and division. Each cell elongated exponentially until its length reached *L*_max_ = 2*L*_0_ + 2*R*, at which point it was replaced by two daughter cells with the length *L*_0_. The doubling time can be calculated to be 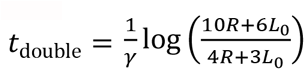. The cell-to-cell and cell-to-substrate contact mechanics were described by linear elastic Hertzian contact mechanics^55^, with a single contact stiffness *E*_0_; note that *E*_0_ corresponds to the modulus of the soft exopolysaccharide in the matrix (∼ 10^2^ Pa) rather than the cell itself, which is much stiffer (∼ 10^5^ Pa). Correspondingly, the *R* value we used (0.8 µm) is larger than the physical size of a cell (∼ 0.4 µm). The parameter values we used were calibrated by rheological measurement and microscopy analysis, and have been shown to successfully capture the dynamics of biofilm-dwelling cells in prior work^33^. The cell-to-substrate adhesion energy was assumed to be linear with the contact area, with adhesion energy density Γ_0_. We incorporated two viscous forces to represent the motion of biofilm-dwelling cells at low Reynold’s number: 1) a bulk viscous drag for all degrees of freedom, and 2) a much larger in-plane surface drag for cells near the substrate, representing the resistance to sliding when a cell is adhered to the substrate via the surface adhesion proteins RbmC/Bap1. The two damping forces also ensured that the cell dynamics were always in the overdamped regime.

We treated the confining hydrogel as a homogenous, isotropic, and linear elastic material using a coarse-grained approach. The geometry of the coarse-grained gel particles was assumed to be spherical with a radius *R*_gel_. The interaction between particles was modeled using a harmonic pairwise potential and a three-body potential related to bond angles. The contact repulsions between the gel particles and the cells as well as between the gel particles and the substrate were described using linear elastic Hertzian contact mechanics. We treated the adhesion between the gel and the substrate using a generalized JKR contact model^56^ and we also included a small viscous damping force to ensure the dynamics remained overdamped. The hydrogel was initialized by annealing the system to achieve an amorphous configuration.

Simulations were initialized with a single cell lying parallel to the substrate and surrounded by gel particles. Initially, a small hemispherical space surrounding the cell was vacated to avoid overlap between the cell and the hydrogel particles. We fixed a small number of hydrogel particles near the boundaries to provide anchoring for the elastic deformation of the hydrogel; however, the boundaries were kept sufficiently far away from the biofilm to minimize any boundary effects. We applied Verlet integration and Richardson integration to numerically integrate the equations of motion for the translational and rotational degrees of freedom, respectively. We implemented the model based on the framework of LAMMPS^57^, utilizing its built-in parallel computing capability. For a more detailed description on the ABS, see Supplementary Information Section 2.

### Quasi-2D agent-based simulations

To further verify the alignment dynamics of the continuum model quantitatively (Eq. 2, Main Text), we developed a set of quasi-2D simulations to mimic the laser irradiation experiments. To simplify the system, the translational and rotational degrees of freedom related to the vertical direction were ignored, while all other parameters were kept the same as the 3D simulations. In each simulation, the bacteria first proliferate normally for 12 hrs, at which point the growth rate of the cells within a radius *r*_0_ from the center of the biofilm was set to 0, mimicking the zone of dead cells caused by laser irradiation (Extended Data Fig. 7). In agreement with experiments, the simulated biofilm was initially randomly oriented (*S* 0); however, cells tended toward an aster pattern and *S* increased over time when the growth void was introduced. The predicted rate at which the cells were driven towards this pattern, in the Lagrangian frame of reference of the cells, is 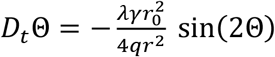. We tested this relationship in the simulation data by comparing the instantaneous angular velocity *D*_t_Θ and 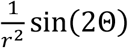 (Fig. 3d). Note that we nondimensionalized the *x*-axis by the final colony radius 25 μm. We varied the radius of the growth void *r*_0_ and repeated the same procedure and for each simulation run, we plotted the slope of the line of best fit versus 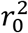 (Fig. 3d inset).

### Data and materials availability

Matlab codes for single-cell segmentation are available online at Github: https://github.com/Haoran-Lu/Segmentation_3D-processing/releases/tag/v1.0. Other data are available upon request.

## Acknowledgments

We thank Drs. A. Mashruwala and Y. Xu for their help in the initial experiments. We thank Drs S. Mao, T. Cohen, and J.-S. Tai for helpful discussions and B. Reed and M. Zhao for help with developing the ABSs.

## Author contributions

J.N. and J.Y. designed and performed the experiments. J.N., Q.Z., H.L., and J.Y. analyzed data. C.L. and S.Z. developed the agent-based simulations. J.N. developed the continuum theory. J.N., C.L., S.Z., and J.Y. wrote the paper.

## Supplementary Materials

Supplementary information is available for this paper.

## References

1. Lecuit, T. & Le Goff, L. Orchestrating size and shape during morphogenesis. Nature 450, 189–192 (2007).

2. Irvine, K. D. & Shraiman, B. I. Mechanical control of growth: ideas, facts and challenges. Development 144, 4238–4248 (2017).

3. Hong, L. et al. Heterogeneity and robustness in plant morphogenesis: from cells to organs. Annu. Rev. Plant Biol. 69, 469–495 (2018).

4. Flemming, H.-C. & Wingender, J. The biofilm matrix. Nat. Rev. Microbiol. 8, 623–633 (2010).

5. Dragoš, A. & Kovács, Á. T. The peculiar functions of the bacterial extracellular matrix. Trends Microbiol. 25, 257–266 (2017).

6. Hall-Stoodley, L., Costerton, J. W. & Stoodley, P. Bacterial biofilms: from the natural environment to infectious diseases. Nat. Rev. Microbiol. 2, 95–108 (2004).

7. Ghannoum, M., Parsek, M., Whiteley, M. & Mukherjee, P. Microbial Biofilms. (ASM Press, 2015).

8. Wuertz, S., Bishop, P. & Wilderer, P. Biofilms in wastewater treatment: an interdisciplinary approach. (IWA Publishing, 2003).

9. Franks, A. E., Malvankar, N. & Nevin, K. P. Bacterial biofilms: the powerhouse of a microbial fuel cell. Biofuels 1, 589–604 (2010).

10. Asally, M. et al. Localized cell death focuses mechanical forces during 3D patterning in a biofilm. Proc. Natl. Acad. Sci. USA 109, 18891–18896 (2012).

11. Trejo, M. et al. Elasticity and wrinkled morphology of Bacillus subtilis pellicles. Proc. Natl. Acad. Sci. USA 110, 2011–2016 (2013).

12. Yan, J. et al. Mechanical instability and interfacial energy drive biofilm morphogenesis. eLife 8, e43920 (2019).

13. Drescher, K. et al. Architectural transitions in Vibrio cholerae biofilms at single-cell resolution. Proc. Natl. Acad. Sci. USA 113, E2066–2072 (2016).

14. Yan, J., Sharo, A. G., Stone, H. A., Wingreen, N. S. & Bassler, B. L. Vibrio cholerae biofilm growth program and architecture revealed by single-cell live imaging. Proc. Natl. Acad. Sci. USA 113, E5337–E5343 (2016).

15. Hartmann, R. et al. Emergence of three-dimensional order and structure in growing biofilms. Nat. Phys. 15, 251–256 (2019).

16. Qin, B. et al. Cell position fates and collective fountain flow in bacterial biofilms revealed by light-sheet microscopy. Science 369, 71–77 (2020).

17. Doostmohammadi, A., Thampi, S. P. & Yeomans, J. M. Defect-mediated morphologies in growing cell colonies. Phys. Rev. Lett. 117, 048102 (2016).

18. You, Z., Pearce, D. J. G., Sengupta, A. & Giomi, L. Geometry and mechanics of microdomains in growing bacterial colonies. Phys. Rev. X 8, 031065 (2018).

19. Dell’Arciprete, D. et al. A growing bacterial colony in two dimensions as an active nematic. Nat. Commun. 9, 4190 (2018).

20. Vernita, D. G., Megan, D.-F., Kristin, K. & Christopher, A. R. Biofilms and mechanics: a review of experimental techniques and findings. J. Phys. D 50, 223002 (2017).

21. Marchetti, M. C. et al. Hydrodynamics of soft active matter. Rev. Mod. Phys. 85, 1143–1189 (2013).

22. Liu, S., Shankar, S., Marchetti, M. C. & Wu, Y. Viscoelastic control of spatiotemporal order in bacterial active matter. Nature 590, 80–84 (2021).

23. He, B., Doubrovinski, K., Polyakov, O. & Wieschaus, E. Apical constriction drives tissue-scale hydrodynamic flow to mediate cell elongation. Nature 508, 392–396 (2014).

24. Keller, P. J., Schmidt, A. D., Wittbrodt, J. & Stelzer, E. H. K. Reconstruction of zebrafish early embryonic development by scanned light sheet microscopy. Science 322, 1065 (2008).

25. Petkova, M. D., Tkacik, G., Bialek, W., Wieschaus, E. F. & Gregor, T. Optimal decoding of cellular identities in a genetic network. Cell 176, 844-855.e15 (2019).

26. Beyhan, S. & Yildiz, F. H. Smooth to rugose phase variation in Vibrio cholerae can be mediated by a single nucleotide change that targets c-di-GMP signalling pathway. Mol. Microbiol. 63, 995–1007 (2007).

27. Andrienko, D. Introduction to liquid crystals. J. Mol. Liq. 267, 520–541 (2018).

28. Fei, C. et al. Nonuniform growth and surface friction determine bacterial biofilm morphology on soft substrates. Proc. Natl. Acad. Sci. USA 117, 7622–7632 (2020).

29. Fong, J. C. N. & Yildiz, F. H. The rbmBCDEF gene cluster modulates development of rugose colony morphology and biofilm formation in Vibrio cholerae. J. Bacteriol. 189, 2319–2330 (2007).

30. Absalon, C., Van Dellen, K. & Watnick, P. I. A communal bacterial adhesin anchors biofilm and bystander cells to surfaces. PLoS Pathog. 7, e1002210 (2011).

31. Berk, V. et al. Molecular architecture and assembly principles of Vibrio cholerae biofilms. Science 337, 236–239 (2012).

32. Fong, J. C. N., Syed, K. A., Klose, K. E. & Yildiz, F. H. Role of Vibrio polysaccharide (vps) genes in VPS production, biofilm formation and Vibrio cholerae pathogenesis. Microbiology 156, 2757–2769 (2010).

33. Beroz, F. et al. Verticalization of bacterial biofilms. Nat. Phys. 14, 954–960 (2018).

34. Başaran, M., Yaman, Y. I., Yuce, T. C., Vetter, R. & Kocabas, A. Large-scale orientational order in bacterial colonies during inward growth. Preprint at https://arxiv.org/abs/2008.05545 (2021).

35. Leslie, F. M. Some constitutive equations for liquid crystals. Arch. Rational Mech. Anal. 28, 265–283 (1968).

36. Grant, M. A. A., Waclaw, B., Allen, R. J. & Cicuta, P. The role of mechanical forces in the planar-to-bulk transition in growing Escherichia coli microcolonies. J. R. Soc. Interface 11, 20140400 (2014).

37. You, Z., Pearce, D. J. G., Sengupta, A. & Giomi, L. Mono-to multilayer transition in growing bacterial colonies. Phys. Rev. Lett. 123, 178001 (2019).

38. Duvernoy, M.-C. et al. Asymmetric adhesion of rod-shaped bacteria controls microcolony morphogenesis. Nat. Commun. 9, 1120 (2018).

39. Marchetti, M. C. et al. Hydrodynamics of soft active matter. Rev. Mod. Phys. 85, 1143–1189 (2013).

40. Doostmohammadi, A., Ignés-Mullol, J., Yeomans, J. M. & Sagués, F. Active nematics. Nat. Commun. 9, 3246 (2018).

41. Beris, Antony. N. & Edwards, B. J. Thermodynamics of flowing systems: with internal microstructure. (Oxford University Press, 1994).

42. Volfson, D., Cookson, S., Hasty, J. & Tsimring, L. S. Biomechanical ordering of dense cell populations. Proc. Natl. Acad. Sci. USA 105, 15346–15351 (2008).

43. You, Z., Pearce, D. J. G. & Giomi, L. Confinement-induced self-organization in growing bacterial colonies. Sci. Adv. 7, eabc8685 (2021).

44. Thompson, D. W. On Growth and Form. (Cambridge University Press, 1992).

45. Copenhagen, K., Alert, R., Wingreen, N. S. & Shaevitz, J. W. Topological defects promote layer formation in Myxococcus xanthus colonies. Nat. Phys. 17, 211–215 (2021).

46. Saw, T. B. et al. Topological defects in epithelia govern cell death and extrusion. Nature 544, 212–216 (2017).

47. Jin, X. & Riedel-Kruse, I. H. Biofilm Lithography enables high-resolution cell patterning via optogenetic adhesin expression. Proc. Natl. Acad. Sci. USA 115, 3698–3703 (2018).

48. Parry, B. R. et al. The bacterial cytoplasm has glass-like properties and is fluidized by metabolic activity. Cell 156, 183–194 (2014).

49. Skorupski, K. & Taylor, R. K. Positive selection vectors for allelic exchange. Gene 169, 47– 52 (1996).

50. Dalia, A. B., McDonough, E. & Camilli, A. Multiplex genome editing by natural transformation. Proc. Natl. Acad. Sci. USA 111, 8937 (2014).

51. Crocker, J. C. & Grier, D. G. Methods of digital video microscopy for colloidal studies. J. Colloid Interface Sci. 179, 298–310 (1996).

52. Wang, Y. et al. Antimicrobial blue light inactivation of pathogenic microbes: State of the art. Drug Resist. Updat. 33–35, 1–22 (2017).

53. Ghosh, P., Mondal, J., Ben-Jacob, E. & Levine, H. Mechanically-driven phase separation in a growing bacterial colony. Proc. Natl. Acad. Sci. USA 112, E2166–E2173 (2015).

54. Lardon, L. A. et al. iDynoMiCS: next-generation individual-based modelling of biofilms. Environ. Microbiol. 13, 2416–2434 (2011).

55. Willis, J. R. Hertzian contact of anisotropic bodies. J. Mech. Phys. Solids 14, 163–176 (1966).

56. Chen, X. & Elliott, J. A. On the scaling law of JKR contact model for coarse-grained cohesive particles. Chem. Eng. Sci. 227, 115906 (2020).

57. Plimpton, S. Fast parallel algorithms for short-range molecular dynamics. J. Comput. Phys. 117, 1–19 (1995).

