## Supplementary Information for "Biofilm self-patterning: mechanical forces drive a reorientation cascade"

### I. Cell segmentation algorithm

After the deconvolved 3D image (Extended Data Fig. 2a, b) was binarized (Extended Data Fig. 2c), a number of different connected components (CCs), each containing multiple cells, were identified. The key difficulty in segmentation lies in the inferior resolution in the *z*-direction in confocal images as well as imperfect rejection of out-of-focus light, which makes the use of a single global threshold inadequate. Therefore, we developed an adaptive local thresholding algorithm for single-cell biofilm image analysis. Specifically, for each CC, we gradually increased the intensity threshold locally and therefore delete voxels whose intensities are below the threshold. After each local binarization step, we checked whether new CCs were generated. For each newly generated CC, if its volume was below a defined volume threshold, we extracted this CC from the image and considered this CC as the core of one individual cell. Information on this individual cell was stored and no further action was performed on this cell until the image-restoration step. If the volume of the CC was still above the volume threshold, we locally increased the threshold until the volume fell below the threshold. After the adaptive local thresholding algorithm, we obtained the cores of all individual cells; however, these cores were smaller than the original cells; they did not represent accurately the location, orientation, and size of the original cells. Therefore, we added a volume restoration step in which the deleted voxels in the segmentation step were merged with the nearest core (Extended Data Fig. 2d, e). By performing the virtual shrinking-expansion step, we maintained the accuracy in both segmentation and in measuring the shape of each bacterium.

### II. Details of the agent-based simulations (ABSs)

<u>*Single cell model*</u>

We model the space occupied by a single cell and its surrounding extracellular matrix as a spherocylinder with length $L$, radius $R$ and therefore volume $V = \frac{4}{3}\pi R^3 + \pi R^2 L$. We assume the cell only grows in length over time while the radius remains the same, where the volume growth obeys the exponential law $\frac{\mathrm{d}V}{\mathrm{d}t} = \gamma V$, with growth rate $\gamma$. We introduce noise during cell growth and division, by assigning different cells with slightly different growth rates, taken from a normal distribution $\gamma \sim \mathrm{N}(\gamma_0, 0.2\gamma_0)$. In the ABSs, cell growth is implemented by increasing the length $L$ by the increment $\Delta L = \gamma \left(\frac{4}{3}R + L\right)\Delta t$, where $\Delta t$ is the timestep.

Cell division is modeled as the instantaneous replacement of a mother cell when it reaches length $L_{\max}$ by two equal sized daughter cells. The slight volume loss after division is ignored to avoid contact overlapping which may cause unphysical reorientations. It follows that the initial length of a daughter cell is $L_0 = \frac{L_{\max}}{2} - R$. Given the growth law, the doubling time can be calculated as $t_{\text{double}} = \frac{1}{\gamma}\log\left(\frac{10R+6L_0}{4R+3L_0}\right)$.

*Cell-cell repulsion*

We only consider the repulsive, elastic contact forces between cells, mediated by the soft exopolysaccharides. Linear elastic Hertzian contact theory is applied to quantify the repulsive contact force on cell $i$ by cell $j$, written as

$$\boldsymbol{F}_{\text{cell-cell},ij} = -\frac{5}{2}E_0R^{1/2}\delta_{ij}^{3/2}\hat{\mathbf{e}}_{ij} \quad \textbf{(1)}$$

where $E_0$ is the effective cell stiffness, $\delta_{ij}$ is the overlapping distance, and $\hat{\mathbf{e}}_{ij}$ is the unit vector parallel to the distance vector $\boldsymbol{d}$. The distance vector is given by connecting two contact points characterizing the smallest distance between two cell centerlines, as shown in Extended Data Fig. 4. The overlapping distance $\delta_{ij}$ is then calculated by

$$\delta_{ij} = 2R - |\boldsymbol{d}|, \quad \textbf{(2)}$$

and contact only occurs if $\delta_{ij} > 0$. Taking the center of mass as the reference point, the moment of contact force is

$$\boldsymbol{M}_{\text{cell-cell},ij} = -\frac{5}{2}s_r\hat{\boldsymbol{n}}_i \times E_0R^{1/2}\delta_{ij}^{3/2}\hat{\mathbf{e}}_{ij} \quad \textbf{(3)}$$

where $\hat{\boldsymbol{n}}_i$ is the cell director and $s_r$ is the parametric coordinate, along the center-line of the cell, of the contact point.

*Cell-to-substrate interactions*

The experimental substrate is modeled as an infinite, two-dimensional rigid plane located at $z = 0$. We again assume linear elastic Hertzian contact mechanics to quantify the repulsive contact between cells and the substrate. We also add an attractive force which depends on the contact area to account for the matrix protein mediated cell-to-surface adhesion.

To formulate the repulsive contact between the cells and the substrate, we apply a generalized Hertzian contact formula that smoothly accounts for the cell orientation dependent contact energy. In this case, the elastic deformation energy is given by $E_{\text{el},i} = E_0R^{1/2}\delta_i^{5/2}$ and the equivalent penetration depth is given by

$$\delta_i^{5/2} = \int_{-L/2}^{L/2}\left[R^{1/2}\left|\hat{\boldsymbol{n}}_{\parallel,i}\right|^2\delta^2(s) + \frac{4}{3}\left(1 - \left|\hat{\boldsymbol{n}}_{\parallel,i}\right|^2\right)\delta^{3/2}(s)\right]\mathrm{d}s \quad \textbf{(4)}$$

where $\hat{\boldsymbol{n}}_{\parallel,i}$ is the normalized projection of the $i$th cell director onto the substrate. The overlap function $\delta(s)$ denotes the overlapping distance between the cell and the substrate at the local cell-body coordinate $-L/2 \le s \le L/2$. Then, the net force $\boldsymbol{F}_{\text{el},i}$ and moment $\boldsymbol{M}_{\text{el},i}$ from the cell-substrate elastic repulsion can be given by

$$\boldsymbol{F}_{\mathrm{el},i} = 2E_0R^{1/2}\int_{L/2}^{-L/2} \hat{\boldsymbol{z}} \cdot \left[R^{1/2}\left|\hat{\boldsymbol{n}}_{\parallel,i}\right|^2\delta(s) + \left(1 - \left|\hat{\boldsymbol{n}}_{\parallel,i}\right|^2\right)\delta^{1/2}(s)\right]\mathrm{d}s \tag{5}$$

$$\boldsymbol{M}_{\mathrm{el},i} = 2E_0R^{1/2}\int_{L/2}^{-L/2} [s\hat{\boldsymbol{n}}_i \times \hat{\boldsymbol{z}}]\left[R^{1/2}\left|\hat{\boldsymbol{n}}_{\parallel,i}\right|^2\delta(s) + \left(1 - \left|\hat{\boldsymbol{n}}_{\parallel,i}\right|^2\right)\delta^{1/2}(s)\right]\mathrm{d}s \tag{6}$$

where $\hat{\boldsymbol{z}}$ is the unit vector perpendicular to the substrate.

We take the adhesion energy between the cells and the substrate to be of the form $E_{\mathrm{ad},i} = -\Sigma_0 A_i$, where $\Sigma_0$ is the adhesion energy density and $A_i$ is the equivalent contact area. The equivalent contact area is given by

$$A_i = \int_{-L/2}^{L/2} a(s)\mathrm{d}s = \int_{-L/2}^{L/2}\left[R^{1/2}\left|\hat{\boldsymbol{n}}_{\parallel,i}\right|^2\delta^{1/2}(s) + \pi R\left(1 - \left|\hat{\boldsymbol{n}}_{\parallel,i}\right|^2\right)\mathrm{H}\big(\delta(s)\big)\right]\mathrm{d}s \tag{7}$$

where $\mathrm{H}(\cdot)$ is the Heaviside step function. The net adhesive force $\boldsymbol{F}_{\mathrm{ad},i}$ and moment $\boldsymbol{M}_{\mathrm{ad},i}$ are:

$$\boldsymbol{F}_{\mathrm{ad},i} = -\Sigma_0\int_{-L/2}^{L/2} \hat{\boldsymbol{z}}\left[\frac{1}{2}R^{1/2}\left|\hat{\boldsymbol{n}}_{\parallel,i}\right|^2\delta^{-1/2}(s)\right]\mathrm{d}s - \hat{\boldsymbol{z}}\Sigma_0\pi R\left(1 - \left|\hat{\boldsymbol{n}}_{\parallel,i}\right|^2\right) \tag{8}$$

$$\begin{aligned}\boldsymbol{M}_{\mathrm{ad},i} = -\Sigma_0\int_{-L/2}^{L/2} [s\hat{\boldsymbol{n}}_i \times \hat{\boldsymbol{z}}]\left[\frac{1}{2}R^{1/2}\left|\hat{\boldsymbol{n}}_{\parallel,i}\right|^2\delta^{-1/2}(s)\right]\mathrm{d}s \\ - [s_0\hat{\boldsymbol{n}} \times \hat{\boldsymbol{z}}]\Sigma_0\pi R\left(1 - \left|\hat{\boldsymbol{n}}_{\parallel,i}\right|^2\right)\end{aligned} \tag{9}$$

where $s_0$ denotes the cell-body coordinate such that $\delta(s_0) = 0$.

*Viscosity*

We consider two sources of viscosity: a bulk viscous force due to the extracellular matrix environment and a surface viscous force due to the substrate. The environmental viscous force and moment are given by Stoke's law,

$$\boldsymbol{F}_{\mathrm{stokes},i} = -\eta_0\boldsymbol{u}_i \tag{10}$$

$$\boldsymbol{M}_{\mathrm{stokes},i} = -\eta_0\int_{L/2}^{L/2} s\hat{\boldsymbol{n}}_i \times (\boldsymbol{\omega}_i \times s\hat{\boldsymbol{n}}_i)\mathrm{d}s = -\frac{\eta_0}{12}\boldsymbol{\omega}_i L^3 \tag{11}$$

where $\eta_0$ is the environmental viscosity, $\boldsymbol{u}$ is the velocity of the center of mass, and $\boldsymbol{\omega}$ is the angular velocity. The substrate viscous force and moment are taken to be of the form

$$\boldsymbol{F}_{\mathrm{surface},i} = -\int_{-L/2}^{L/2}\frac{\eta_1 a(s)}{R}[\boldsymbol{u}_i(s) - (\boldsymbol{u}_i(s)\cdot\hat{\boldsymbol{z}})\hat{\boldsymbol{z}}]\mathrm{d}s \tag{12}$$

$$\boldsymbol{M}_{\mathrm{surface},i} = -\int_{-L/2}^{L/2}\frac{\eta_1 a(s)}{R}s\hat{\boldsymbol{n}}_i \times [\boldsymbol{u}_i(s) - (\boldsymbol{u}_i(s)\cdot\hat{\mathbf{z}})\hat{\boldsymbol{z}}]\mathrm{d}s \tag{13}$$

where $\eta_1$ is the viscous coefficient along the substrate.

*Equations of motion*

The equations of motion for each cell are given by Newton's rigid body dynamics:

$$\begin{pmatrix} \boldsymbol{F}_{\mathrm{net},i} \\ \boldsymbol{M}_{\mathrm{net},i} \end{pmatrix} = \begin{bmatrix} m & \mathbf{0} \\ \mathbf{0} & \mathbf{I}_i \end{bmatrix} \begin{pmatrix} \dot{\boldsymbol{u}}_i \\ \dot{\boldsymbol{\omega}}_i \end{pmatrix} + \begin{pmatrix} \mathbf{0} \\ \boldsymbol{\omega}_i \times \mathbf{I}_i \boldsymbol{\omega}_i \end{pmatrix} \quad \textbf{(14)}$$

where $\boldsymbol{F}_{\mathrm{net},i}$ and $\boldsymbol{M}_{\mathrm{net},i}$ are the total force and moment vector, and $\mathbf{I}$ is the moment of inertia. All of the variables are expressed in the body-fixed coordinate system, then transformed into the global coordinate system. We add a small random noise to the net force and moment vectors of the cells at every timestep ($10^{-7}E_0R^2$ for forces and $10^{-7}E_0R^3$ for moments).

*Choice of parameters for cellular dynamics*

The list of physical constants, which were used to successfully capture the verticalization instability in biofilm-dwelling cells in prior work[1], are used here and given in Table S2.

*Geometry of coarse-grained gel system*

We model the agarose gel using a coarse-grained particle-based approach. We treat the gel as a collection of spherical particles, each with radius $R_{\mathrm{gel}}$. We treat the interactions of the spheres using a spring network model to recapitulate the elastic behavior of the hydrogel. The pairwise interaction energy between the gel particles is $E_{\mathrm{gel},2} = \Sigma_{ij} \frac{k_r}{2}\left(\xi_{ij} - \xi_0\right)^2$, where $\xi_{ij}$ is the distance between particle $i$ and $j$, $\xi_0$ is the equilibrium distance, and $k_r$ is the spring constant. Furthermore, to impart a shear modulus to the system, we also include a three-body interaction energy where $E_{\mathrm{gel},3} = \Sigma_{ijk} \frac{k_\zeta}{2}\left(\zeta_{ijk} - \zeta_0\right)^2$, where $\zeta_{ijk}$ is the bond angle formed by particle $i$, $j$, and $k$.

*Interactions between gel particles and cells*

We again apply linear elastic Hertzian contact theory to describe the repulsive interaction between the coarse-grained gel particles and the cells. Similar to the contact between cells, the elastic contact energy can be written as $E_{\mathrm{gel-cell},ij} = E_1\left(\frac{1}{R} + \frac{1}{R_{\mathrm{gel}}}\right)^{-1/2} \delta_{ij}^{5/2}$, where $E_1$ is the contact stiffness between gel and cell, and $\delta_{ij}$ is the overlapping distance between the $i$th gel particle and $j$th cell. The contact stiffness $E_1$ can be given by the relation $\frac{1}{E_1} = \frac{1-\mu_{\mathrm{gel}}^2}{Y_{\mathrm{gel}}} + \frac{1-\mu_{\mathrm{cell}}^2}{Y_{\mathrm{cell}}}$, where $Y$ and $\mu$ are Young's modulus and Poisson's ratio, respectively.

*Interactions between the gel particles and the substrate*

There exist two types of interactions between the coarse-grained gel particles and the substrate. First, the gel particles make elastic contact with the substrate, again described by Hertzian contact theory for the contact between a sphere and a flat surface, where the elastic contact energy is given by $E_{\mathrm{gel-surface},i} = E_1 R_{\mathrm{gel}}^{1/2} \delta_i^{5/2}$. Second, we introduce adhesion between the gel particles and substrate which provides an energy barrier to delamination; we take the energy to be of the form $E_{\mathrm{ad,gel},i} = -\Sigma_1 A_{\mathrm{gel},i}$, where $\Sigma_1$ is the adhesion energy density. The equivalent contact area is given by $A_{\mathrm{gel},i} = \pi R_{\mathrm{gel}} \delta_i$, where $\delta_i$ is the overlap between the gel particle and the substrate.

*Equation of motion*

We only consider the three translational degrees of freedom for gel particles, neglecting the rotational degrees of freedom. Therefore, the equations of motion are given by Newton's second law $\mathbf{F}_{\text{tot},i} = m\boldsymbol{a}_i$, where $\boldsymbol{F}_{\text{tot},i}$ is the net force and $\boldsymbol{a}_i$ is the acceleration. To prepare the initial amorphous stress-free geometry, we begin with a body-centered cubic crystalline geometry with lattice parameter $a$, where $a = 1.3R_{\text{gel}}$. Subsequently, we assigned the system with an initial temperature of $300K$ and annealed it until it reached a final configuration that is amorphous and stress-free.

*Choice of parameters*

The spring constant $k_r$ and equilibrium length $\xi_0$: The Young's modulus of the coarse-grained gel system is given by $Y = \frac{1}{V}\frac{\partial^2(E_{\text{gel},2}+E_{\text{gel},3})}{\partial\epsilon^2} \simeq \frac{k_r}{2\xi_0}^2$, under the condition $k_\zeta \ll k_r$. Generally, a smaller $\xi_0$ leads to a denser gel system and better approximation to a continuum solid. Here we choose $\xi_0 = 0.6$ μm as a result of a trade-off between simulation quality and computational cost, as the simulation time is proportional to $\frac{1}{{\xi_0}^3}$. This $\xi_0$ value leads to $k_r = 6 \times 10^{-3}\text{Nm}^{-1}$, corresponding to the experimentally measured $Y = 5kPa$.

Radius of coarse-grained gel particle $R_{\text{gel}}$: In order to mimic the continuum constraints posed by the hydrogel in the experiment, the coarse-grained system should not have significant defects larger than the volume of a single cell, which means that $R_{\text{gel}}$ should be larger than $\xi_0$. On the other hand, $R_{\text{gel}}$ cannot be significantly larger than the cell radius $R$, as this will introduce unphysical contacts at the biofilm-gel interface. Taking both requirements into consideration, we choose $R_{\text{gel}} = 1.0$ μm, which is nearly double the equilibrium distance $\xi_0 = 0.6$ μm and we keep $R_{\text{gel}} \approx R$.

Cell-gel contact modulus $E_1$: Hertzian contact mechanics gives the equivalent contact modulus between two elastic bodies by $\frac{1}{E_1} = \frac{1-\mu_{cell}^2}{Y_{cell}} + \frac{1-\mu_{gel}^2}{Y_{gel}}$, where $Y$ is the Young's modulus and $\mu$ is Possion's ratio of the two contacting elastic bodies. Given the bulk rheology measurement $Y_{\text{cell}} = 500$ Pa, $\mu_{\text{cell}} = \mu_{\text{gel}} = 0.49$, and $Y_{\text{gel}} = 5$ kPa[3,4], it follows $E_1 = 600$ Pa. In the simulations, we choose $E_1 = 1500$ Pa, somewhat larger than $E_0$, to avoid the unphysical situation in which cells penetrate into the gel during growth.

The three-body interaction parameter $k_\zeta$ and equilibrium angle $\zeta_0$: To make the amorphous coarse-grained gel system stable, we choose $\zeta_0 = 120°$ to force the system to deviate from its original cubic configurations. We select $k_\zeta$ such that we attain incompressible solid behavior in the gel $G = Y/3$ , where $G$ is the shear modulus and $Y$ is Young's modulus.

These parameters are listed in Table S3.

### III. Single-cell surface adhesion stability analysis

Previous studies have described how cells in 2D bacterial colonies can be peeled from the substrate through a buckling-like instability[5–8]; however, the reason why biofilm-dwelling cells self-organize into a verticalized core, that is, remain attached but oriented away from the substrate, remains to be shown. In this section, we consider the stability of surface-adhered cells in the presence of compression from neighboring cells to determine why surface adhesion facilitates stably surface-anchored, verticalized cells. Specifically, we consider a minimal model consisting of three spherocylinders following the same physics described in the agent-based modelling section above; however, we limit our analysis below to cells confined to the $xz$ plane, without any external gel confinement, and only experiencing the effects of the bulk viscosity for analytical tractability. We assume that the cells are not growing and instead mimic the effects of growth-induced compression by bringing the outer cells closer together. All three cells initially start out parallel to the substrate in close contact with each other, and the two outer cells are brought closer to each other to squeeze the central cell (see the schematic in Figure 2f). We find that the middle cell evolves through four phases upon increasing compression: **I.** the cell remains horizontal and the elastic potential energy in the cell increases; **II.** the middle cell rotates from horizontal to vertical and all potential energy is consequently released; **III.** the middle cell remains vertical, and the elastic potential energy increases again when the two outer cells touch the center cell; and **IV.** the middle cell becomes unstable and is ejected from the substrate. The transition from I to II corresponds to a "verticalization" instability[1], from II to III corresponds to a trivial geometric transition, and from III to IV corresponds to a "pinch-off" instability. Note that the pinch-off instability is locally (marginally) stable, but under sufficiently large $z$-perturbations, the middle cell gets ejected from the substrate. Experimentally, we expect that many factors could contribute to the perturbations including but not limited to: deviation of the cell shape from a perfect spherocylinder, fluctuating protein bonds, and variations in the adhesion protein concentration, etc. Importantly, we find that for the parameters listed in Table S2, the critical energy corresponding to the verticalization instability is much smaller than that of the pinch-off instability, leading to preferential verticalization of horizontal cells rather than pinch-off of verticalized cells (Fig. 2f). In what follows below, we consider these two instabilities in more detail paying specific attention to the role of the cell-to-surface adhesion.

The equations of motion for the position $\boldsymbol{r_i} = (x_i, z_i)$, and director $\hat{\boldsymbol{n}}_i = (n_x, n_z)$ of cell $i$ written using Lagrangian mechanics, are:

$$\eta_0 l \dot{\boldsymbol{r}}_i = -\frac{\partial \sum E_i}{\partial \boldsymbol{r_i}},$$
$$\frac{\eta_0 l^3 \dot{\hat{\boldsymbol{n}}}_i}{12} = -\frac{\partial \sum E_i}{\partial \hat{\boldsymbol{n}}_i} + \lambda \frac{\partial F_i}{\partial \hat{\boldsymbol{n}}_i}, \qquad \textbf{(15A,B)}$$

where $\lambda$ is a Lagrange multiplier corresponding to the constraint $F_i = |\hat{\boldsymbol{n}}_i| - 1 = 0$. Here $\sum E_i$ is the total potential energy of cell $i$ and includes cell-to-cell repulsion, as well as cell-to-surface adhesion and repulsion terms.

In phase I, in the absence of any compression from neighboring cells, the balance of surface adhesion and repulsion leads to an equilibrium cell configuration $\hat{\boldsymbol{n}} = (1,0)$ at an apparent height

$$z_{0,h} = R\left(1 - \left(\frac{\tilde{A}}{4}\right)^{\frac{2}{3}}\right), \qquad \textbf{(16)}$$

where $\tilde{A} = \Sigma_0 / R E_0$ is the dimensionless adhesion coefficient. In the presence of compression, we consider perturbations away from this stable point. Taking $x$ as the distance between the centers of the two outer cells (we assume that the outer cells are at the same equilibrium height), the elastic energy in the cell of interest due to cell-to-cell contacts is $E_{\mathrm{cc}} = 2E_0 R^{1/2}(2R - d)^{5/2}$ where the shortest distance between the cells is $d = L + 2R - x/2$. Combining with the cell-surface adhesion and cell-surface repulsion terms, we find the evolution equation for small $n_z$ is given by

$$\dot{n}_z = \Omega(x) n_z, \quad \textbf{(17)}$$

where the sign of $\Omega$ determines the stability of the fixed-point. Extended Data Figure 6a shows plots of $\Omega$ for different dimensionless adhesion coefficients. For small values of $\tilde{A}$, which are most physically relevant, the zeros of $\Omega$ are only weakly dependent on $\tilde{A}$. This analysis suggests that the verticalization instability only weakly depends on adhesion, that is, the energy barrier to verticalization is nearly constant.

In phase III, again assuming no compression from neighboring cells, the balance of surface adhesion and repulsion leads to an equilibrium verticalized configuration $\hat{\boldsymbol{n}} = (0,1)$ with height

$$z_{0,v} - \frac{L}{2} = R\left(1 - \left(\frac{3\pi\tilde{A}}{4}\right)^{2/3}\right). \quad \textbf{(18)}$$

Note that the effective penetration depth (the degree to which the soft cell is flattened against the substrate) is larger for verticalized cells than for horizontal cells. Again, we consider compression due to two outer horizontal cells (at a height $z_{0,h}$). Here we look for finite-sized perturbations away from the fixed point; the evolution equation of the height of the verticalized cell is given by

$$\dot{z} = w(x, z). \quad \textbf{(19)}$$

We look for points $(x, z)$ such that $w(x, z) > 0$. We only consider points $0 \leq \frac{L}{2} + R - z < R - z_{0,h}$ as points $z - \frac{L}{2} < z_{0,h}$ results in no net vertical force such that the central cell remains stably attached to the substrate. Specifically, we focus on points very close to $z = \mathrm{L}/2 + R$ to determine the critical energy required to produce marginally unstable cells. Extended Data Figure 6b shows plots of $w(x, L/2 + 0.999R)$ for different adhesion coefficients. Contrary to the verticalization transition, in this case we find a strong dependence of the zeros of $w$ on $\tilde{A}$; therefore, the energy barrier to pinch-off depends strongly on cell-to-surface adhesion.

We combine these two stability analyses into a single plot of the energy landscape experienced by the center cell for different adhesion energies (Extended Data Figure 6c). We see that for small adhesion energies, the difference in the two energy barriers is small, whereas for large adhesion energies, the pinch-off instability requires orders of magnitude more energy than the verticalization instability. This energy difference between the two instabilities leads to more stably adhered verticalized cells. This analysis explains the observed positive correlation between the adhesion energy and the fraction of stably verticalized cells (Extended Data Fig. 3, 4): adhesion energy affects the stability of cells with respect to the pinch-off instability but much less so to the verticalization instability.

### IV. Continuum model for growth-induced macroscopic cell ordering

In this section, we present a minimal coarse-grained description to explain the growth and self-organization observed in the basal layer of *V. cholerae* biofilms. We first consider the simpler case with one population of cells that either generate in-plane growth or not, before considering the complete two-phase model which accounts for pressure-dependent cell verticalization and out-of-plane growth.

*Theoretical framework*

We limit our consideration to the basal plane of the biofilm and assume it is a growing, quasi-2D system with *mesoscopic* nematic order tensor $\boldsymbol{Q} = 2\langle\hat{\boldsymbol{n}}\otimes\hat{\boldsymbol{n}} - \boldsymbol{I}/2\rangle$[9]. Based on its symmetries, in 2D, $\boldsymbol{Q}$ can also be written as

$$\boldsymbol{Q} = q\begin{bmatrix}\cos(2\theta) & \sin(2\theta)\\ \sin(2\theta) & -\cos(2\theta)\end{bmatrix} \tag{20}$$

where $\theta$ is the angle of the average head-less director and $q$ is the scalar order parameter, which quantifies the degree to which the cells are locally ordered. In the analysis below we will assume that this scalar order parameter is constant.

We assume the bacterial cells grow with a uniform growth rate $\gamma$ but that the growth can be out-of-plane, resulting in a spatially varying in-plane growth rate $g(\boldsymbol{r})$. Taking the cell density as $c$, and the coarse-grained growth-induced velocity field as $\boldsymbol{u}$, mass conservation requires

$$\partial_t c + \nabla\cdot(c\boldsymbol{u}) = gc. \tag{21}$$

The density is nearly uniform in the experiment, that is, $c(\boldsymbol{r}) = c_0 + \epsilon c_1(\boldsymbol{r})$, where $\epsilon \ll 1$. To leading order, Eq. (21) becomes

$$\nabla\cdot\boldsymbol{u} = g. \tag{22}$$

The growth-induced velocity field is found by integrating the spatially dependent growth rate. In this analysis, we will neglect density fluctuations $c_1$, but it can be related to the fluctuation in local pressure due to varying local configurations.

The evolution of the nematic order tensor is described by the Beris-Edward equation[10]

$$(\partial_t + \boldsymbol{u}\cdot\nabla)\boldsymbol{Q} = \lambda\boldsymbol{E} - (\boldsymbol{\omega}\cdot\boldsymbol{Q} - \boldsymbol{Q}\cdot\boldsymbol{\omega}) + \Gamma\boldsymbol{H}, \tag{23}$$

where $\lambda$ is the flow-alignment parameter and $\lambda > 1$ for rod-shaped objects that tend to align in shear. Here $\boldsymbol{E} = \frac{1}{2}(\nabla\boldsymbol{u} + \nabla\boldsymbol{u}^T - (\nabla\cdot\boldsymbol{u})\boldsymbol{I})$ and $\boldsymbol{\omega} = \frac{1}{2}(\nabla\boldsymbol{u} - \nabla\boldsymbol{u}^T)$ are the traceless strain-rate and vorticity tensors and $\boldsymbol{H}$ is the molecular field. $\Gamma\boldsymbol{H}$ relaxes $\boldsymbol{Q}$ towards a bulk state with minimal spatial variation in the director. However, we do not expect $\Gamma\boldsymbol{H}$ to play an important role in biofilm ordering because biofilm-dwelling cells secrete exopolysaccharides which are soft and deformable and act as a "cushion" between cells. This can be seen in the biofilm from the Δ*BC* mutant (Fig. 1e) in which minimal local alignment was observed. Contrast this with the Δ*vpsL* mutant and other bacterial colonies[11–13] where there is significant local alignment (Fig. 1f) when no exopolysaccharides are present. Finally, force balance requires

$$\nabla\cdot\boldsymbol{\Pi} = \eta\boldsymbol{u}, \tag{24}$$

where $\boldsymbol{\Pi}$ is the stress tensor in the biofilm, whose divergence is balanced by surface friction. Here we have assumed that energy dissipation is dominated by surface friction rather than viscous dissipation inside the biofilm, corresponding to the so-called "dry" limit[14]. We take the frictional force density to be linearly proportional to the velocity and the friction coefficient $\eta$ to be isotropic and constant. In many active nematic systems, the active stress is anisotropic and dependent on $\boldsymbol{Q}$. For instance, in motile cell colonies, cells generate anisotropic active stresses[15–17] while in growing bacterial colonies a number of different constitutive relations have been proposed[11,12,18,19]. Although at the microscopic scale, these forces may be anisotropic, on the mesoscopic scale pertaining to our theory, they are expected to be isotropic on average. This leads to the assumption of an isotropic stress field which is balanced by surface friction,

$$\nabla p = \eta \boldsymbol{u}. \tag{25}$$

Pressure in this system arises due to compression from neighboring cells, mediated by the exopolysaccharides, which arises from cell proliferation.

To close the model, we impose the following set of boundary and initial conditions. By symmetry, we expect that the velocity goes to zero at the center of the biofilm,

$$\boldsymbol{u}(\boldsymbol{0}, t) = \boldsymbol{0}, \tag{26}$$

that is, the center of the biofilm does not move and

$$\frac{\partial \boldsymbol{Q}}{\partial r}(\boldsymbol{0}, t) = \boldsymbol{0}. \tag{27}$$

We also assume that the pressure accumulates towards the center of the biofilm and that at the outer edge of the biofilm the pressure is zero,

$$p(r = r_e, t) = 0, \tag{28}$$

where $r_e$ denotes the edge of the biofilm. The outer radius evolves through the kinematic condition

$$\frac{dr_e}{dt} = u_r(r_e). \tag{29}$$

Finally, we assume that the orientation of the biofilm is described by some initial order parameter tensor

$$\boldsymbol{Q}(r, \psi, t = 0) = \boldsymbol{Q}_{\boldsymbol{0}}(r, \psi), \tag{30}$$

which can also be written as $\theta(r, \psi, t = 0) = \theta_0(r, \psi)$.

Equations (**22**, (**23**, (**25** and boundary conditions and initial conditions Eq. (**26**)-(**30**) form the basis of the theoretical model. In the following subsections, we consider a few idealized cases to determine under which conditions biofilms self-organize into an aster pattern. Throughout, we will also make the simplifying assumption that all variables except for $\boldsymbol{Q}$ are axisymmetric and that there is no azimuthal flow, $\boldsymbol{u} = u_r \hat{r}$.

<u>Uniform growth limit</u>

If all growth is in-plane, then $g = \gamma$ is uniform and constant. Integrating Eq. (**22** we find the velocity field is

$$u_r = \frac{\gamma r}{2}, \tag{31}$$

and the strain-rate and vorticity tensors are exactly zero, $\boldsymbol{E} = \boldsymbol{\omega} = \boldsymbol{0}$. Solving Eq. (**23** yields

$$\boldsymbol{Q}(r, \psi, t) = \boldsymbol{Q}_0\left(re^{-\frac{\gamma t}{2}}, \psi\right). \tag{32}$$

*In this case, $\boldsymbol{Q}$ is simply stretched by growth, and there is no tendency for $\boldsymbol{Q}$ to rotate or align in any given direction*. An initially isotropic system will remain isotropic as it grows (Extended Data

Fig. 7a-c)[12,20]. This is what happens in the 2D expansion of bacterial colonies, which remain macroscopically isotropic as they grow. Although microscopic forces can align cells locally, they cannot introduce any long-range order. Note that in two dimensions axisymmetric growth is a special case where there is no net strain in the system: if instead the system were confined into a rectangular channel, where growth is unidirectional, the strain rate tensor would not be zero and therefore there would be net alignment along the direction of the channel[18,19,21,22].

*Alignment due to a growth void*

Now suppose there is a radius $r_0$ inside which the cells have verticalized and are angled out of plane. These cells do not contribute to the growth of the basal plane and so

$$g(r) = \begin{cases} 0 \text{ if } r < r_0 \\ \gamma \text{ if } r \geq r_0 \end{cases}. \quad \textbf{(33)}$$

Substituting into Eq. (**22**, the growth-induced velocity is

$$u_r = \begin{cases} 0 & \text{if } r < r_0 \\ \gamma(r - r_0^2/r)/2 & \text{if } r \geq r_0 \end{cases}. \quad \textbf{(34)}$$

The corresponding strain rate and vorticity tensors in polar coordinates ($\boldsymbol{E_p}, \boldsymbol{\omega_p}$) are

$$\begin{aligned} \boldsymbol{E_p} &= \boldsymbol{0}, & \boldsymbol{\omega_p} &= \boldsymbol{0} \text{ if } r < r_0 \\ \boldsymbol{E_p} &= \begin{bmatrix} \left(\frac{\partial u_r}{\partial r}\right) - \frac{\gamma}{2} & 0 \\ 0 & \frac{u_r}{r} - \frac{\gamma}{2} \end{bmatrix}, & \boldsymbol{\omega_p} &= \boldsymbol{0} \text{ if } r \geq r_0. \end{aligned} \quad \textbf{(35)}$$

Under uniform growth $\partial u_r/\partial r$ and $u_r/r$ are equal to $\gamma/2$ , leading to no net deviatoric strain-rate. In contrast, in the presence of a growth void, these two terms become unequal, leading to a net deviatoric strain-rate. This is because the radial gradient in velocity is no longer balanced by azimuthal stretching due to outward expansion. For a velocity of the form given in Eq. (**34**, the strain rate and vorticity tensors in polar coordinates are

$$\begin{aligned} \boldsymbol{E_p} &= \boldsymbol{0}, & \boldsymbol{\omega} &= \boldsymbol{0} \text{ if } r < r_0 \\ \boldsymbol{E_p} &= \frac{\gamma r_0^2}{2r^2}\begin{bmatrix} 1 & 0 \\ 0 & -1 \end{bmatrix}, & \boldsymbol{\omega} &= \boldsymbol{0} \text{ if } r \geq r_0. \end{aligned} \quad \textbf{(36)}$$

Next, the nematic order parameter $\boldsymbol{Q}$ can be transformed to a polar coordinate system, given by

$$\boldsymbol{Q_p} = \boldsymbol{R^T Q R} = q\begin{bmatrix} \cos(2(\theta - \psi)) & \sin(2(\theta - \psi)) \\ \sin(2(\theta - \psi)) & -\cos(2(\theta - \psi)) \end{bmatrix}, \quad \textbf{(37)}$$

where $\psi$ is the azimuthal angle and $\boldsymbol{R} = \begin{bmatrix} \cos\psi & -sin\,\psi \\ sin\,\psi & \cos\psi \end{bmatrix}$ is the coordinate transformation matrix from cartesian to polar coordinates.

Finally, given $\boldsymbol{E_p}$, $\boldsymbol{\omega_p}$ and $\boldsymbol{Q_p}$, the matrix form of Eq. **(23)** in polar coordinates is

$$q\left(\partial_t + u_r\frac{\partial}{\partial r}\right)\begin{bmatrix} \cos(2(\theta - \psi)) & \sin(2(\theta - \psi)) \\ \sin(2(\theta - \psi)) & -\cos(2(\theta - \psi)) \end{bmatrix} = \frac{\lambda\gamma r_0^2}{2r^2}\begin{bmatrix} 1 & 0 \\ 0 & -1 \end{bmatrix}, \quad \textbf{(38)}$$

for $r \geq r_0$, which yields two scalar equations

$$-\sin(2(\theta-\psi))\left(\frac{\partial\theta}{\partial t}+u_r\frac{\partial\theta}{\partial r}\right)=\frac{\lambda\gamma r_0^2}{4qr^2}, \qquad \textbf{(39A,B)}$$
$$\cos(2(\theta-\psi))\left(\frac{\partial\theta}{\partial t}+u_r\frac{\partial\theta}{\partial r}\right)=0.$$

Defining $\Theta = (\theta - \psi)$ and combining the two scalar equations above, we find that for $r \geq r_0$,

$$\partial_t\Theta + u_r\partial_r\Theta = -\frac{\lambda\gamma r_0^2}{4qr^2}\sin(2\Theta). \qquad \textbf{(40)}$$

By first ignoring the advective term, we see that $\partial_t\Theta \sim -\sin(2\Theta)$ which has stable fixed points at $\Theta = n\pi$ or $\theta = \psi + n\pi$, characteristic of an aster pattern. *In other words, the rightmost term, which arises due to gradients in the growth-induced velocity field, drives cells to reorient towards an aster pattern.* The same observation was made during inward growth in bacterial colonies where deviation of the velocity field from that of isotropic growth resulted in radial alignment of cells[20].

More generally, if we suppose that instead of a growth void, there is a general differential growth rate: $\gamma$ for $r < r_0(t)$ and $\gamma + \Delta\gamma$ for $r \geq r_0(t)$ then the evolution equation for $\Theta$ for $r > r_0$ is given by

$$\partial_t\Theta + u_r\partial_r\Theta = -\frac{\lambda\Delta\gamma r_0^2}{4qr^2}\sin(2\Theta). \qquad \textbf{(41)}$$

If the biofilm has a faster growing outer rim $\Delta\gamma > 0$, the bacteria will tend to organize into an aster pattern. If, on the other hand, the biofilm has a faster growing core, i.e. $\Delta\gamma < 0$, $\Theta$ will have stable fixed points at $\Theta = n\pi + \pi/2$, characteristic of a vortex pattern. This means that biofilms that exhibit excess growth in the core will be driven to form a vortex.

Returning to Eq. **(40**, we solve it by the method of characteristics, assuming some initial maximum biofilm radius $r_e$, which yields

$$\cot[\Theta(r,\psi)] = \cot[\Theta_0(r',\psi)]\exp\left[\frac{\lambda}{2q}\left(\gamma t + \log\left(\frac{r'^2}{r^2}\right)\right)\right], \qquad \textbf{(42)}$$

where $r' = \left((r^2 - r_0^2)e^{-\gamma t} + r_0^2\right)^{1/2}$ and $\Theta_0 = \theta_0 - \psi$ corresponds to the angular representation $\boldsymbol{Q}_0$. Plots of Eq. **(42** are given in Extended Data Figures 7d-f. Note that, although cells tend to align in an aster pattern, the aligning torque becomes weaker as cells are advected outwards. Therefore, $S$ does not necessarily reach a value of 1 (Extended Data Fig. 7g). Depending on the initial size of the growth void as well as the flow alignment parameter, different degrees of radial alignment are reached in the final state, with a larger $r_0/r_e$ leading to overall more radial alignment.

To be more general, now instead of a finite void, suppose there is an arbitrary velocity field $u_r(r,t)$ corresponding to an arbitrary growth rate $g(r,t)$. Following the same procedure as above, we find the evolution equation for $\Theta$ is

$$\partial_t\Theta + u_r\partial_r\Theta = -f(r,t)\sin(2\Theta), \qquad \textbf{(43)}$$

where $f = (\lambda r/4q)\partial_r(u_r/r)$ quantifies the aligning torque due to gradients in the flow field. Equation **(43)** thus describes a generalized time-evolution equation for the orientation field due to gradients in the flow velocity.

*Two-phase active nematic model for cell verticalization and alignment*

We expand the active nematic model in the previous subsection to include the verticalization dynamics in a two-phase model consisting of vertical and horizontal cells. Consider a quasi-2D layer of cells on a substrate where the cells are either parallel (horizontal) or tilted (vertical) with respect to the substrate and let $\rho$ denote the fraction ($0 \leq \rho \leq 1$) of vertical cells and $1-\rho$ therefore denote the fraction of horizontal cells. Horizontal cells proliferate and generate their progeny in the basal layer of interest, while vertical cells do not contribute to the expansion to the basal layer. The growth-induced stress generated by the horizontal cells also cause them to verticalize with a pressure dependent rate $C(p)$. The evolution equation for the fraction of these two populations of cells is therefore given by:

$$\begin{gathered}\partial_t \rho + \nabla \cdot (\boldsymbol{u}\rho) = C(p)(1-\rho) \\ \partial_t(1-\rho) + \nabla \cdot \big(\boldsymbol{u}(1-\rho)\big) = \gamma(1-\rho) - C(p)(1-\rho).\end{gathered} \qquad \textbf{(44A,B)}$$

Given that cell verticalization requires a minimum threshold pressure $p_t$, we make the simple assumption that the conversion rate is a linear function of the excess pressure

$$C(p) = \beta \frac{p - p_t}{p_t} H(p - p_t), \qquad \textbf{(45)}$$

where $H$ denotes the Heaviside function. Combining Eq. (**44**A and (**44**B yields

$$\nabla \cdot \boldsymbol{u} = \gamma(1-\rho). \qquad \textbf{(46)}$$

We initiate the biofilm with some initial radius $r_{e,0}$ and assume all cells are horizontal to begin with, $\rho(r, t=0) = 0$. Finally, we assume symmetry at the origin which requires $\frac{\partial \rho}{\partial r} = 0$ and that $\rho$ is axisymmetric. Equations (**25**, (**43)**, (**44**, (**45**, (**46** with the above mentioned initial and boundary conditions make up the full two-phase continuum model for cellular ordering driven by verticalization.

*Non-dimensionalization*

We non-dimensionalize Eq. (**25**, (**43)**, (**44**, (**45**, (**46** by the following dimensional scales: time $t_d = 1/\gamma$, pressure $p_d = p_t$, length $r_d = (p_t/\eta\gamma)^{1/2}$ and velocity $u_{r,d} = r_d\gamma$, which yields

$$\frac{\partial \tilde{p}}{\partial \tilde{r}} = \tilde{u}_r, \qquad \textbf{(47)}$$

$$\frac{\partial \Theta}{\partial \tilde{t}} + u_{\tilde{r}} \frac{\partial \Theta}{\partial \tilde{r}} = -\tilde{L}\tilde{r} \frac{\partial \tilde{u}_r/\tilde{r}}{\partial \tilde{r}} \sin(2\Theta), \qquad \textbf{(48)}$$

$$\frac{\partial \rho}{\partial \tilde{t}} + \frac{1}{r}\frac{\partial(\tilde{r}\tilde{u}_r\rho)}{\partial \tilde{r}} = \tilde{\beta}(1-\rho)(p-1)H(p-1), \qquad \textbf{(49)}$$

$$\frac{1}{r}\frac{\partial(\tilde{r}\tilde{u}_r)}{\partial \tilde{r}} = 1 - \rho. \qquad \textbf{(50)}$$

with boundary conditions $p(\tilde{r}_e) = 0$, $\tilde{u}_r(0) = 0$, $d_t\tilde{r}_e = \tilde{u}_r(\tilde{r}_e)$, $\partial_{\tilde{r}}\Theta|_{\tilde{r}=0} = 0$, and $\partial_{\tilde{r}}\rho|_{\tilde{r}=0} = 0$ and initial conditions $\Theta(\tilde{r},\psi,0) = \Theta_0(\tilde{r},\psi)$ and $\rho(\tilde{r},0) = 0$. At the boundary, when solving for $\Theta$, we impose a sharp velocity gradient $\Delta\tilde{u}_r/\Delta\tilde{r} \approx -\tilde{u}_r(\tilde{r}_e)/(\tilde{l}_{\text{Cell}}/2)$ where $\tilde{l}_{\text{Cell}}$ is the average length of a cell, to account for the fact that across the boundary cells there is a sharp velocity gradient which tends to align cells tangentially to the boundary[7,12]. There are two key parameters that control the evolution of the system: the dimensionless flow alignment parameter $\tilde{L} = \lambda/4q$ and the dimensionless verticalization rate $\tilde{\beta} = \beta/\gamma$. The flow alignment parameter dictates how quickly the cells will align to a straining flow, while the verticalization rate dictates how quickly horizontal cells are converted to vertical cells. For simplicity, we assume that the only distinguishing feature of the nonadherent mutant is that $\beta = 0$; however, we would also expect that in the absence of adhesion, $\eta$ would also be smaller and lead to an underestimation of the characteristic length-scale.

*Choice of parameters*

We choose the following set of parameters when solving for the evolution of the system.

Growth rate $\gamma$: The number of bacteria in a three-dimensional biofilm was measured over time and the growth rate was found to be $\gamma \approx 0.6$ hr.$^{-1}$

Threshold verticalization pressure $p_t$: From the theoretical verticalization analysis, using experimentally calibrated parameters, it was found that the critical cell overlap distance at which point verticalization happens is $\delta \sim 0.125$ µm. From this value, the critical cell-cell contact pressure can be calculated as $p_d = 4E\delta^{\frac{1}{2}}/(3\pi R^{*1/2})$ where $E$ is the contact modulus of the cells which we estimate to be $500$ Pa using bulk rheological measurements (shear modulus of a bulk WT* biofilm was measured to be $\sim 200$ Pa) and $R^* = R/2 = 0.4$ µm. This yields a threshold pressure of $p_t \sim 130$ Pa.

Friction coefficient $\eta$: Using the surface viscosity $\eta_1$ defined in Table S2, we estimate the friction coefficient as $\eta = \eta_1/A$ where $A$ is the surface footprint of the cell which we estimate to be $A \sim 3$ µm$^2$. The friction coefficient is therefore $\eta \sim 7 \times 10^4$ Pa s/µm$^2$.

Characteristic length scale $r_d$: The characteristic length-scale $r_d = (p_t/\eta\gamma)^{1/2}$ in the system corresponds to the characteristic size of biofilm when verticalization begins. For the parameters defined above, this length scale is $r_d \sim 3.4$ µm.

Mean cell length $l_{\text{Cell}}$: Taking the average of the largest and smallest cylindrical sections of the spherocylindrical cells (corresponding to right before and right after division) gives a length of $l_{\text{Cell}} = 1.8$ µm.

Initial radius of biofilm $r_{e,0}$: We initialize the biofilm to have an area equivalent to the footprint of a single cell. This gives a value $r_{e,0} \sim 1$ µm.

We solve the dimensionless Eq. (**47**(**50** using a finite difference scheme with explicit time-stepping, and up-winding of advective terms. Example solutions are given in Extended Data Figure 10. We fit the two unknown dimensionless parameters to the experimental results and find that $\tilde{\beta} \approx 2.5$ and $\tilde{L} \approx 1.5$. These parameters compare favorably to what has been measured before. Namely, Beroz *et al.*[1] measured in ABS that in a biofilm growing without a confining agarose gel, $\beta \sim 2.5$ hr.$^{-1}$. Here we find a somewhat smaller value, $\beta \sim 1.5$ hr.$^{-1}$, likely owing to the fact that confinement due to the agarose gel tends to impart normal stresses that suppress verticalization. Nonetheless, we find that the results are only weakly dependent on $\beta$ for $\beta > \gamma$ (Fig. 4a-d; Extended Data Fig. 10d-k). Finally, we estimate $q$ by measuring the cell-cell orientation correlation for neighboring cells in the basal layer of the experimental biofilms and find $q \sim 0.5$ and so $\lambda \sim 3$. Measuring the intrinsic flow alignment parameter is difficult as it is dependent on many factors including the size, shape, aspect ratio and local nematic order[23], however, a value of 3 is in reasonable agreement with values of 0.3-2 reported for other growing and non-growing bacterial systems[11,18,24].

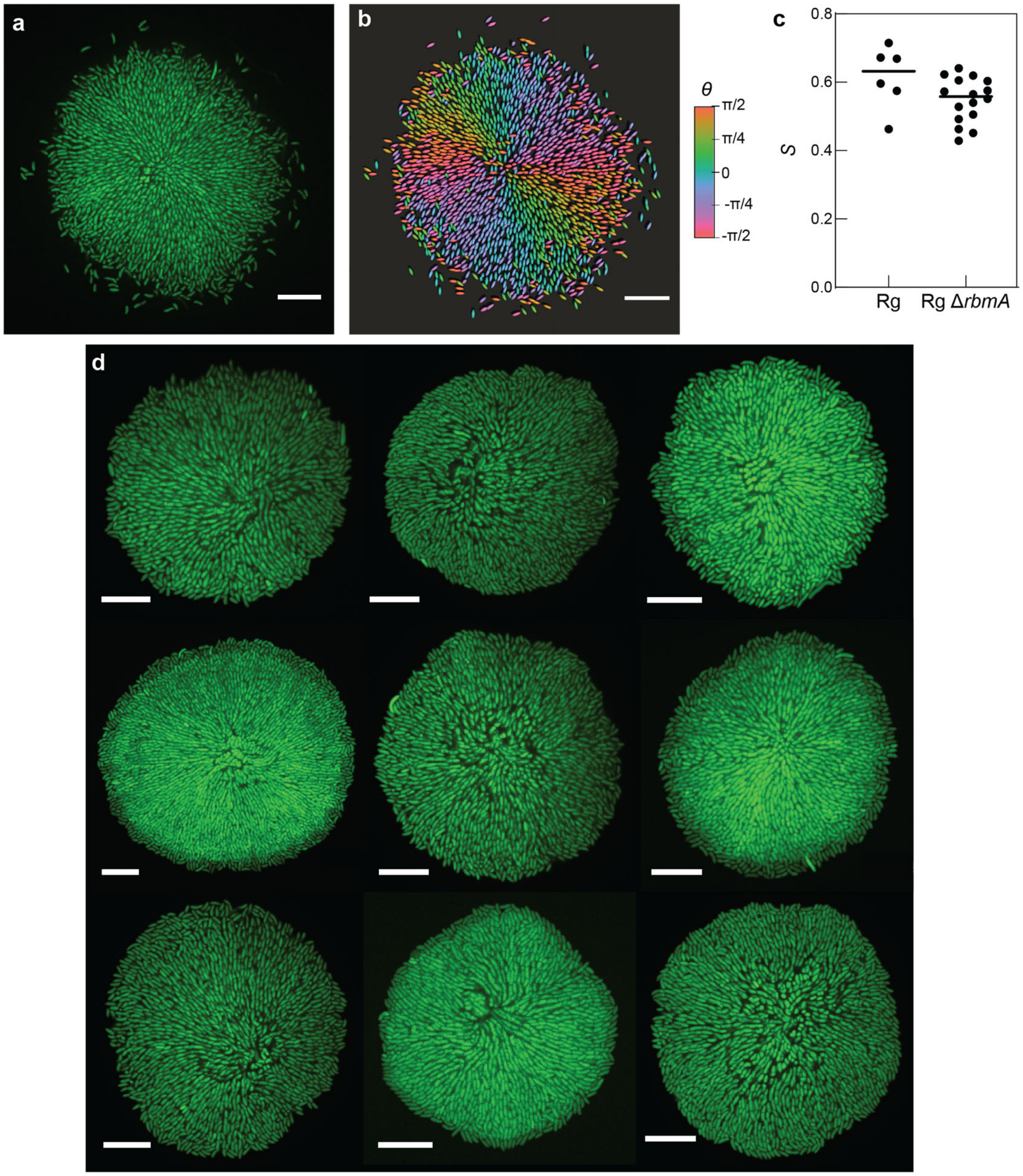

**Extended Data Fig. 1| *V. cholerae* biofilms robustly and reproducibly form aster patterns. a**, Cross-sectional view of the basal plane of a rugose biofilm with an intact *rbmA* gene. **b**, The same biofilm reconstructed where cells are colored based on the angle of the in-plane director. **c**, Radial order parameter *S* in biofilms formed by the rugose wild-type and RgΔ*rbmA* strain (labeled as WT* in the main text). We find no statistical difference between the two strains (unpaired *t*-test with Welch's correction) under this growth condition, suggesting that the presence of the cell-to-cell adhesion protein does not interfere with the global organization mechanism. In fact, the rugose wild-type biofilms display the same radial organization even when growing freely on a solid substrate without confinement by the agarose gel, an observation that has been reported[25,26] but never explained. Therefore, we believe that the same macroscopic ordering mechanism applies generally to biofilms grown in any geometry. In this work, we used a confined geometry to increase the basal area of the biofilm, in order to focus on this emergent organization. **d,** Nine different mature Rg∆*rbmA* biofilms all exhibiting the aster pattern. The core of the aster, typically defined by the location of the founder cell, does not necessarily coincide with the geometric center of the biofilm due to stochasticity during early biofilm development. Scale bars, 10 µm.

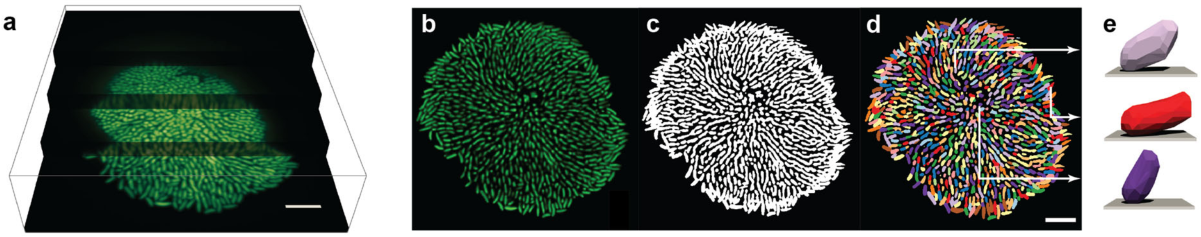

**Extended Data Fig. 2| Cell segmentation procedure. a,** Representative raw 3D image of a WT* biofilm. **b-d**, Intermediate steps during biofilm segmentation; after deconvolution with a measured point spread function (**b**), after binarization (**c**), after segmentation using an adaptive local thresholding algorithm (**d**). **e**, 3D reconstruction of three different segmented cells at three different locations in the biofilm. Scale bars, 10 μm.

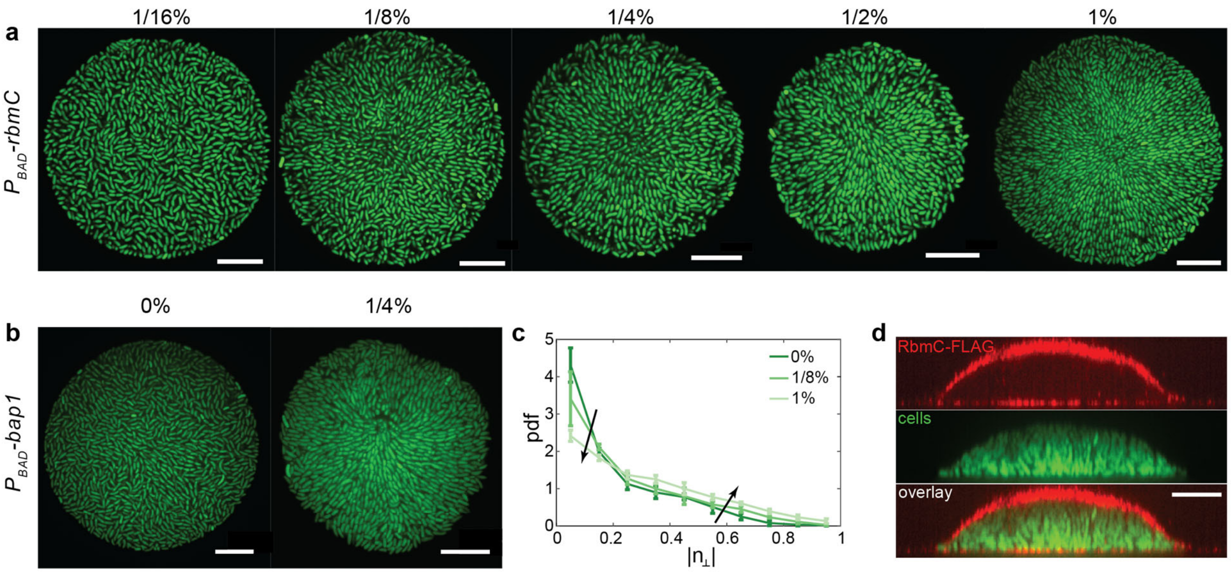

**Extended Data Fig. 3| Cell-to-surface adhesion controls macroscopic cell organization. a**, Cross-sectional view of the basal layer of biofilms with arabinose-inducible *rbmC* expression. In the presence of different concentrations of arabinose, a transition from disorder-to-order was observed with increasing arabinose. **b**, A similar disorder-to-order transition was seen with arabinose-inducible *bap1* expression. Bap1 and RbmC have been shown to contribute to cell-to-surface adhesion in a partially redundant fashion[27]. **c**, Probability distribution function (pdf) of the degree of verticalization $|n_\perp|$ showed a shift to more verticalized cells and less horizontal cells with higher *rbmC* expression controlled by arabinose concentration ($\geq$ 4 biofilms; error bars correspond to std). **d**, Spatial distribution of RbmC-3×FLAG stained with Cy3-conjugated anti-FLAG antibodies shows that RbmC localizes on both the glass and gel surfaces. Scale bars, 10 μm.

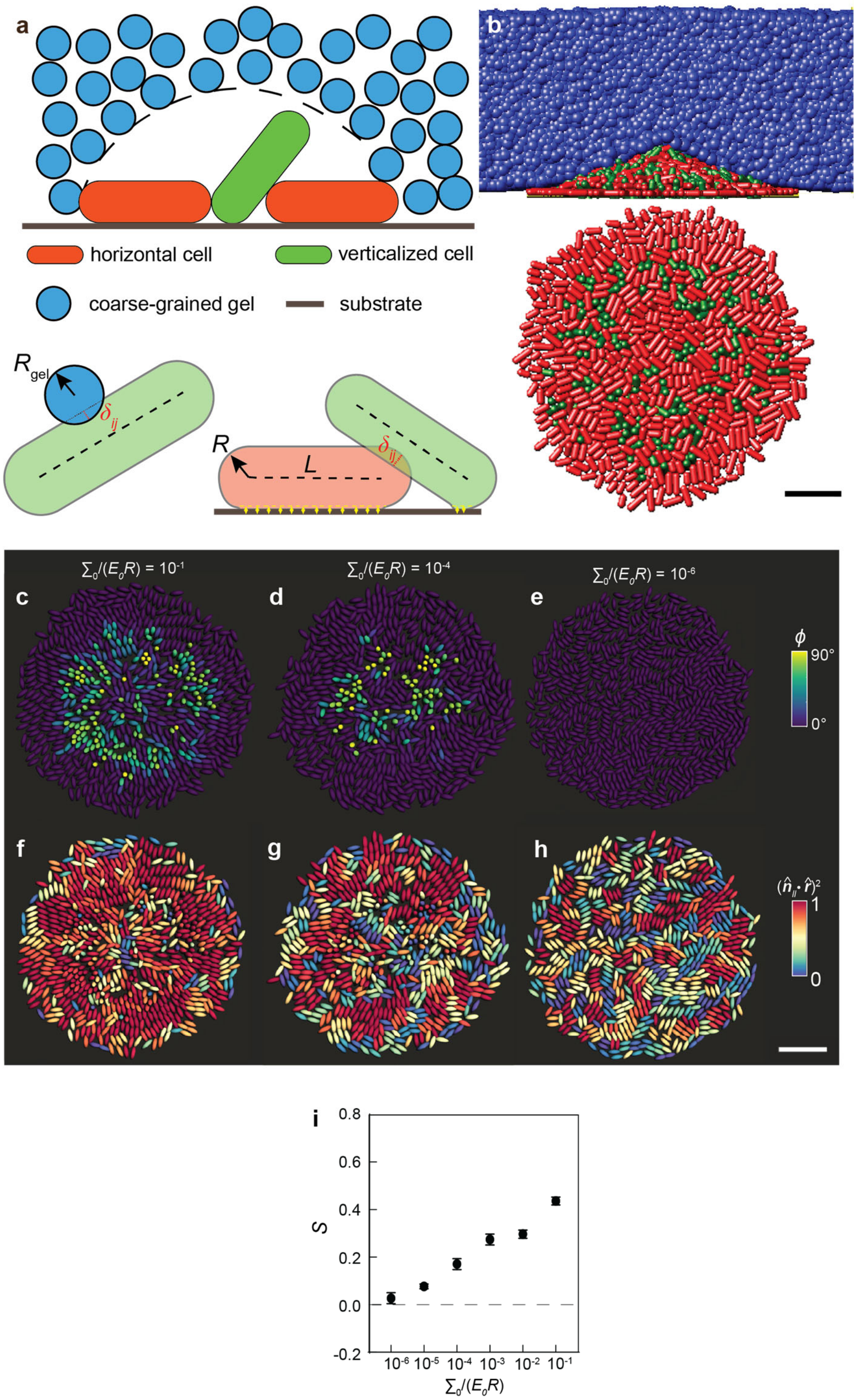

**Extended Data Fig. 4| Agent-based simulations of 3D biofilms with varying adhesion. a**, Schematic illustration of the agent-based simulation consisting of biofilm dwelling cells, which are modeled as growing spherocylinders with length $L(t)$ and radius $R$, and the surrounding hydrogel, which is modeled using a coarse-grained particle system. **b**, Side view (*Top*) and bottom view (*Bottom*) of a representative simulated biofilm. In the bottom view, gel particles are omitted for clarity. Verticalized cells ($n_\perp > 0.5$) are labelled green, and horizontal cells ($n_\perp \leq 0.5$) are labelled red. Scale bar, 5 μm. **c-e**, Cells in the basal layer, color-coded by the angle $\phi$ each cell makes with the substrate, in biofilms possessing diffferent adhesion strengths. The dimensionless adhesion strength $\tilde{A} = \Sigma_0/RE_0$ is $10^{-1}$, $10^{-4}$, and $10^{-6}$ for **c** to **e**, respectively. **f-h**, The same biofilms as **c-e** with cells color-coded by the degree of radial alignment $(\hat{\boldsymbol{n}}_\parallel \cdot \hat{\boldsymbol{r}})^2$. Scale bar, 10 μm. **i,** Averaged $S$ in biofilms with different dimensionless adhesion strengths. Each data point corresponds to the mean and error bars correspond to the standard deviation of 10 different simulated biofilms.

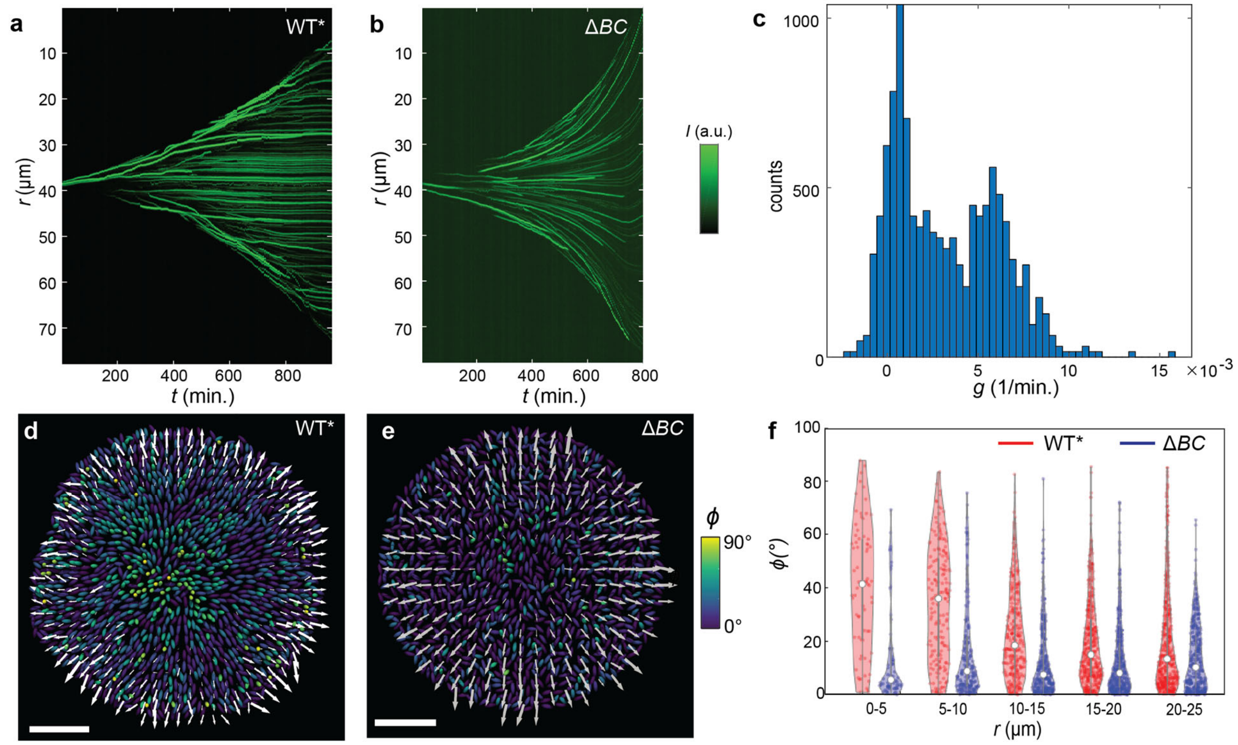

**Extended Data Fig. 5| Biofilms spontaneously develop zones with differential growth due to adhesion-mediated stable verticalization. a, b**, Kymograph of puncta trajectories in a WT* (**a**) and a Δ*BC* mutant biofilm (**b**). In the WT* biofilm a stationary central region develops and expands. In contrast, in the mutant biofilm, no stationary region exists. **c**, Histogram of apparent growth rates showing a bimodal distribution of "growing" and "non-growing" cells. Reconstructed image of a biofilm from a WT* (**d**) and a Δ*BC* mutant biofilm (**e**), where cells are color-coded by the angle they make with respect to the bottom substrate. Overlain are arrows denoting the measured velocity field. **f**, Violin plot showing the distribution of cell orientations at different radii, for the WT* and Δ*BC* mutant biofilms, respectively. The WT* biofilm was able to sustain verticalized cells leading to a growth void in the middle and a nonlinear velocity profile, whereas the Δ*BC* mutant is unable to sustain verticalized cells, therefore leading to a linear velocity profile. Scale bars, 10 µm.

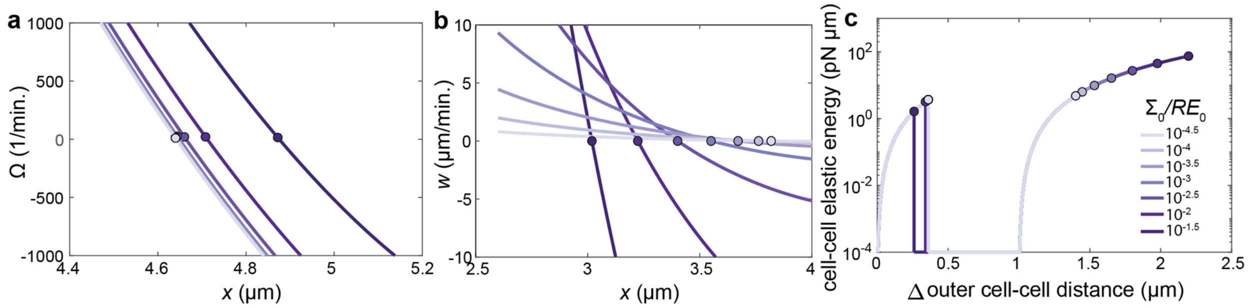

**Extended Data Fig. 6| Comparison of energy landscapes for different adhesion energies. a**, **b**, Determination of the instability point for the verticalization (**a**) and pinch-off instabilities (**b**). Regions in which $\Omega > 0, w > 0$ correspond to regions where fixed points are unstable. Circles denote the transition between stable and unstable behavior. Here $\dot{n}_z = \Omega(x)n_z$ and $\dot{z} = w(x, L/2 + 0.999R)$. **c**, Elastic energy due to cell-to-cell contacts in a cell being squeezed incrementally by two neighbors for different dimensionless adhesion energies, $\widetilde{\mathrm{A}} = \Sigma_0/RE_0$ (see schematic in Fig. 2F). Circles denote the points at which instabilities occur. Here the energy landscape is calculated using the results of the stability analysis.

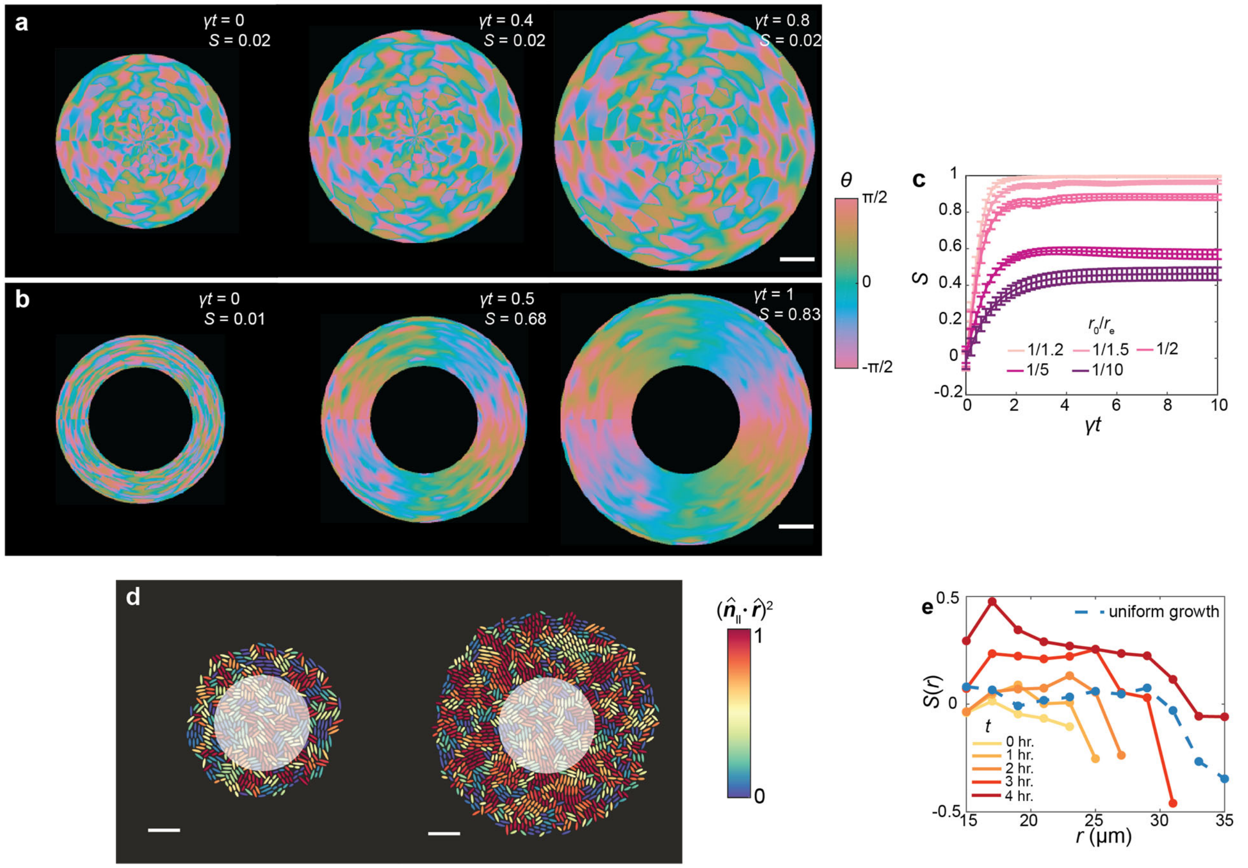

**Extended Data Fig. 7| Continuum modelling and ABSs confirm that a growth void is sufficient to drive radial cell alignment. a**, **b**, Evolution of the director field for a biofilm without (**a**) and with (**b**) a growth void. In panel **b** the radius of the void is $r_0 = (3/5)r_{\mathrm{e}}$, where $r_{\mathrm{e}}$ corresponds to the biofilm size when the growth void is introduced. **c**, Evolution of $S$ averaged over the growing region for different dimensionless initial void sizes $r_0/r_{\mathrm{e}}$. The dimensionless flow alignment parameter $\tilde{L} = \lambda/4q = 1.5$. Each data point corresponds to the mean and error bars correspond to the standard deviation of 10 different random initializations. Radial order emerges over time but plateaus because, unlike the WT* biofilm with an expanding verticalized core, the size of the void is constant in this set of calculations and its effect on cell reorientation is limited to cells near the void due to the $1/r^2$ dependence of the driving force. **d**, Configuration of a 2D biofilm (*left*) right before imposing a growth void and (*right*) 4hr after imposing a growth void (highlighted by the white circle). The color denotes the degree to which the cell orientation coincides with the radial director $(\hat{\boldsymbol{n}}_{\parallel} \cdot \hat{\boldsymbol{r}})^2$. **e,** Azimuthally averaged $S(r)$ in biofilms after the creation of a growth void at different times. Dashed blue line shows the result from a control biofilm without a growth void. Scale bars, 10 μm.

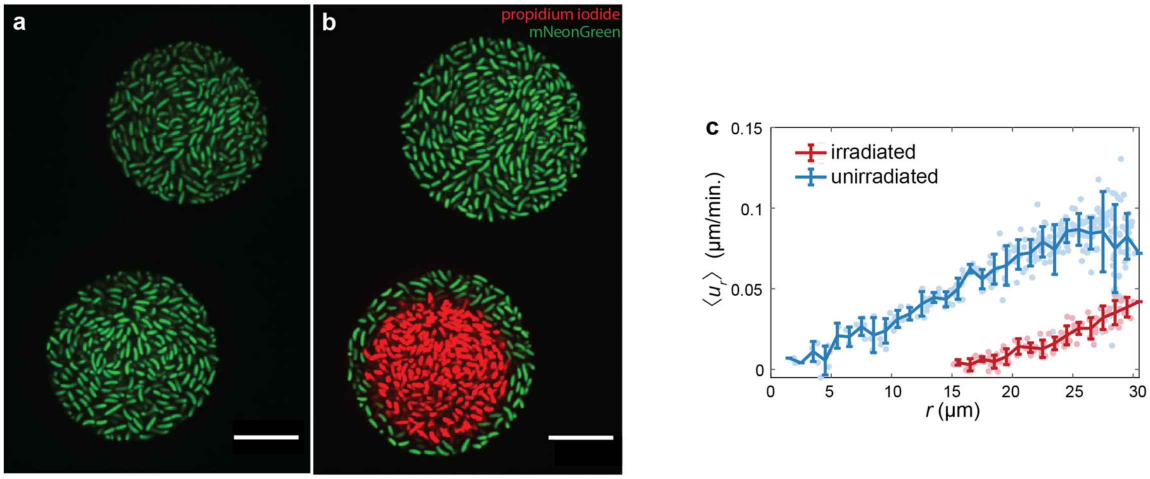

**Extended Data Fig. 8| Laser irradiation causes cell death and results in a growth void. a**, Two $\Delta BC$ biofilms in close proximity before laser irradiation. **b**, The same biofilms after being irradiated in a circular pattern using 405 nm light. Dead cells are stained with propidium iodide. **c**, The measured radial velocity field 1 hour after irradiation (red) and without irradiation (blue). Scale bars, 10 μm.

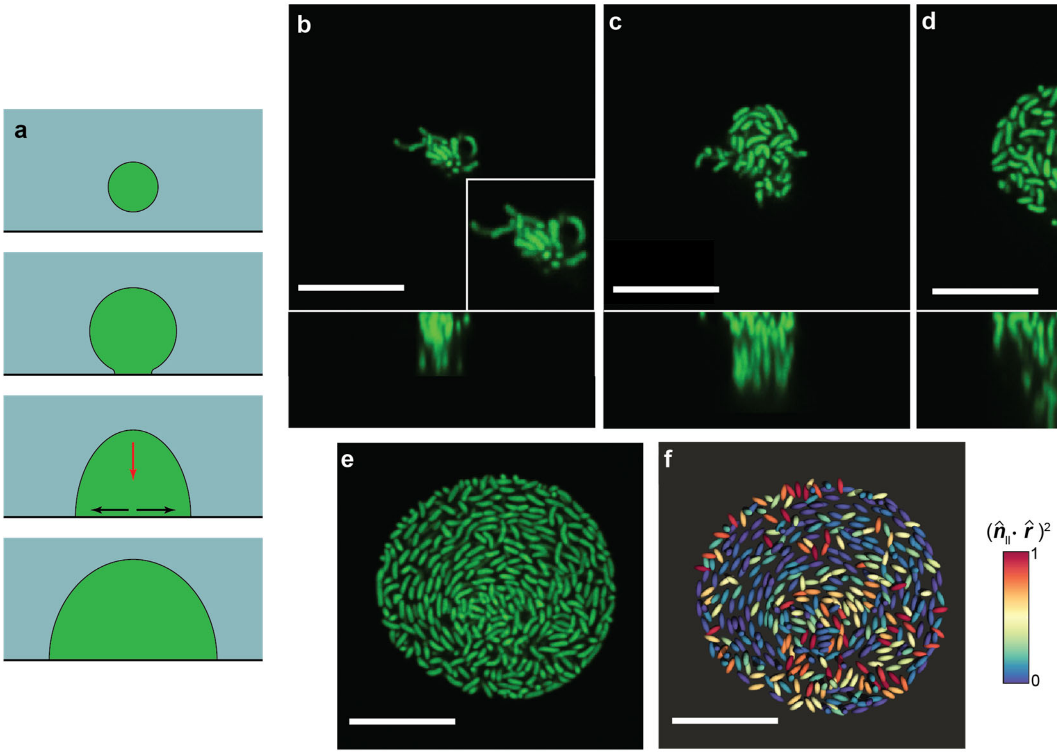

**Extended Data Fig. 9| Nonadherent biofilms have the potential to form vortex structures due to non-uniform growth.** To probe if vortex structures were possible, we imaged $\Delta BC$ biofilms that were initially embedded inside the bulk of the gel instead of starting at the gel-glass interface. This is because, once such a biofilm reached the interface, it preferentially grew at the interface with little increase in biofilm height, leading to excess growth introduced into the basal layer from the bulk, mostly at the center. This process is given schematically in panel **a**. Initially, the biofilm grows completely inside the gel without being in contact with the glass surface. Eventually, the growing biofilm begins to spread between the gel and the glass surfaces and cells grow preferentially along this interface. In this case, excess growth is supplied by cells in the bulk (red arrow) which add to the in-plane growth rate at the center of the basal layer (black arrows). Excess growth at the center gives rise to a negative $f(r,t)$ in Main Text Eq. 2 (see Supplementary Information Section 4 for more discussion), therefore providing a driving force to align cells azimuthally. **b-e**, Time-lapse imaging of a growing $\Delta BC$ biofilm in which a vortex-like structure appears ($t = 6, 9, 12, 15$ hrs). A zoom-in image is provided in **b** when the embedded biofilm first touches the glass surface; at this moment cells are randomly oriented. **f**, Reconstructed biofilm from panel **e** where cells are color-coded based on the degree of radial alignment $(\hat{\boldsymbol{n}}_{\|} \cdot \hat{\boldsymbol{r}})^2$. A value of zero denotes cells that are oriented azimuthally (circumferentially).

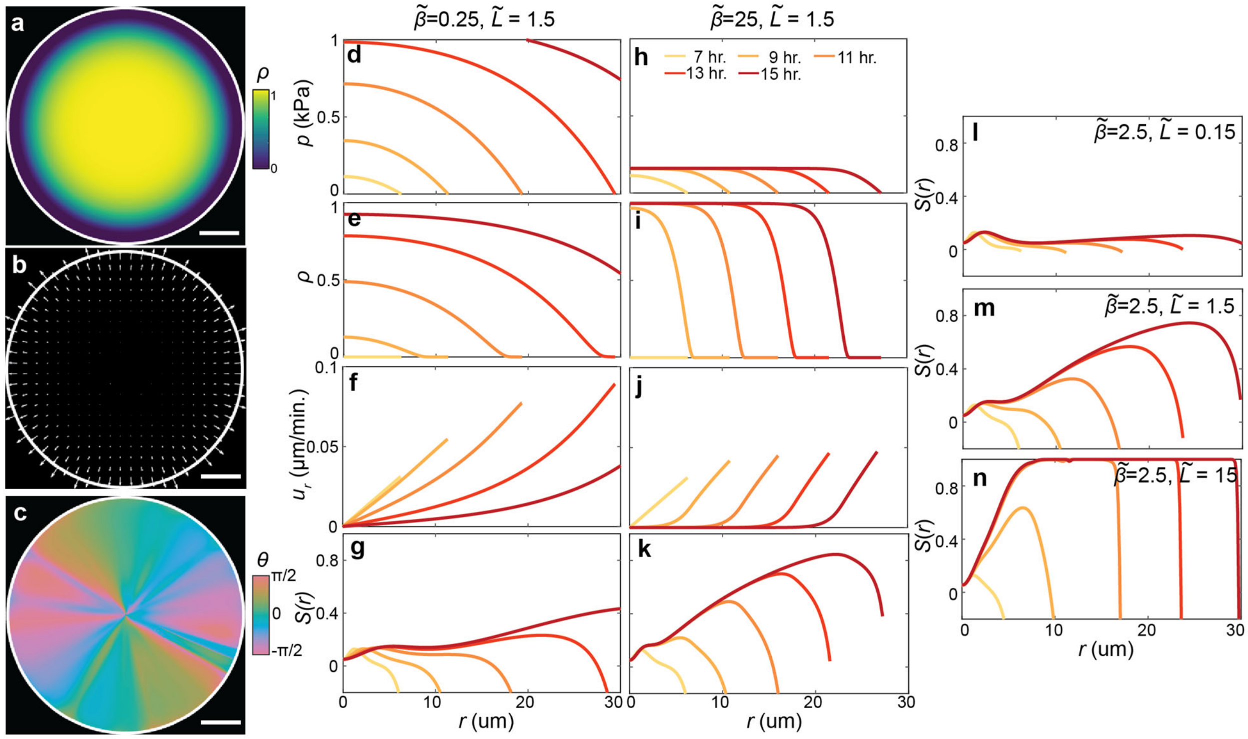

**Extended Data Fig. 10| Representative results from two-phase active nematic model. a-c**, Representative 2D plots of the fraction of verticalized cells $\rho$ (**a**), the velocity field (**b**), and the director field (**c**) for dimensionless verticalization rate $\tilde{\beta} = 2.5$ and dimensionless flow alignment parameter $\tilde{L} = 1.5$. Scale bars, 10 µm. **d-k**, Results of the model for a small verticalization rate $\tilde{\beta} = 0.25$ (**d-g)** and for a large verticalization rate $\tilde{\beta} = 25$ (**h-k)**. **l-n**, Evolution of $S$ for three different flow alignment rates $\tilde{L}$ while keeping the same $\tilde{\beta} = 2.5$. The colors denote model results at different times in panels **d-n**.

**Table S1: List of the strains used in this study**

| Strains | Genotype | Source |
|---|---|---|
| JN007 | *vpvc*$^{W240R}$*, ΔrbmA, Δvc1807::Ptac-mNeonGreen*-Spec$^{R}$ | This study |
| JN009 | *vpvc*$^{W240R}$*, ΔrbmA, Δbap1, ΔrbmC, Δvc1807::Ptac-mNeonGreen*-Spec$^{R}$ | This study |
| JN010 | *vpvc*$^{W240R}$*, ΔvpsL, Δvc1807::Ptac-mNeonGreen*-Spec$^{R}$ | This study |
| JN148 | *vpvc*$^{W240R}$*, ΔrbmA, ΔlacZ::Ptac-mNeonGreen-μNS, Δvc1807::Ptac-mScarletI*-Spec$^{R}$ | This study |
| JN150 | *vpvc*$^{W240R}$*, ΔrbmA, Δbap1, ΔrbmC, ΔlacZ::Ptac-mNeonGreen-μNS, Δvc1807::Ptac-mScarletI*-Spec$^{R}$ | This study |
| JN035 | *vpvc*$^{W240R}$*, ΔrbmA, Δbap1, ΔrbmC, Δvc1807::Ptac-mNeonGreen*-Spec$^{R}$, pJY056 | This study |
| JN036 | *vpvc*$^{W240R}$*, ΔrbmA, Δbap1, ΔrbmC, Δvc1807::Ptac-mNeonGreen*-Spec$^{R}$, pJY057 | This study |
| JY489 | *vpvc*$^{W240R}$*, ΔrbmA, rbmC-3×FLAG, Δvc1807::Ptac-mNeonGreen*-Spec$^{R}$ | This study |
| **Plasmid** | | |
| pJY056 | Kan$^{R}$, *araC-*$P_{BAD}$*-rbmC* | [26] |
| pJY057 | Kan$^{R}$, *araC-*$P_{BAD}$*-bap1* | [26] |

**Table S2: List of cell parameters used in the agent-based simulations.**

| $R(\mu\text{m})$ | $L_{\text{max}}(\mu\text{m})$ | $E_0(\text{Pa})$ | $\gamma(\text{s}^{-1})$ | $\Sigma_0(\text{Nm}^{-1})$ | $\eta_0(\text{Pa}\cdot\text{s})$ | $\eta_1(\text{Pa}\cdot\text{s})$ |
|---|---|---|---|---|---|---|
| 0.8 | 3.6 | 300 | $3.12 \times 10^{-4}$ | $7.3 \times 10^{-6}$ | 20 | $2 \times 10^5$ |

**Table S3: List of gel parameters used in the agent-based simulations.**

| $R_{\mathrm{gel}}(\mu\mathrm{m})$ | $E_1(\mathrm{Pa})$ | $k_{\mathrm{r}}\ (\mathrm{Nm}^{-1})$ | $k_{\zeta}(\mathrm{Nm}^2)$ | $\xi_0(\mu\mathrm{m})$ | $\zeta_0(°)$ | $\Sigma_1(\mathrm{Nm}^{-1})$ |
|---|---|---|---|---|---|---|
| 1.0 | 1500 | $6 \times 10^{-3}$ | $6.6 \times 10^{-4}$ | 0.6 | 120 | $6 \times 10^{-2}$ |

**Movie captions**

**Movie S1:** A time-lapsed cross-sectional view of the basal plane of a growing WT* biofilm. Imaging began immediately after seeding the founder cell. The total duration of the movie is 20 hr. The scale bar is 10 μm.

**Movie S2:** A time-lapsed cross-sectional view of the basal plane of a WT* biofilm with mNeonGreen labelled puncta. Imaging began 8 hr. after seeding the founder cell. The total duration of the movie is 16 hr. The scale bar is 10 μm.

**Movie S3:** Representative results of the agent-based simulations. (*Left*) Bottom view of a biofilm in which cells are colored red if horizontal ($n_{\perp} \leq 0.5$) or green if vertical ($n_{\perp} > 0.5$). (*Right*) Side view of the same biofilm (red/green) and surrounding course-grained gel particles (blue). The duration of the movie is 18 hr.

**Movie S4:** Representative results of a quasi-2D agent-based simulation in which a growth void is introduced at the center of a growing biofilm. The movie is slowed down 1.5-fold after the introduction of the growth void. The total duration of the movie is 16 hr.

### References

1. Beroz, F. *et al.* Verticalization of bacterial biofilms. *Nat. Phys.* **14**, 954–960 (2018).
2. Ostoja-Starzewski, M., Sheng, P. Y. & Alzebdeh, K. Spring network models in elasticity and fracture of composites and polycrystals. *Comput. Mater. Sci.* **7**, 82–93 (1996).
3. Yan, J. *et al.* Mechanical instability and interfacial energy drive biofilm morphogenesis. *eLife* **8**, e43920 (2019).
4. Yan, J. *et al.* Bacterial biofilm material properties enable removal and transfer by capillary peeling. *Adv. Mater.* **30**, 1804153 (2018).
5. Grant, M. A. A., Wacław, B., Allen, R. J. & Cicuta, P. The role of mechanical forces in the planar-to-bulk transition in growing *Escherichia coli* microcolonies. *J. R. Soc. Interface* **11**, 20140400 (2014).
6. You, Z., Pearce, D. J. G., Sengupta, A. & Giomi, L. Mono- to multilayer transition in growing bacterial colonies. *Phys. Rev. Lett.* **123**, 178001 (2019).
7. Su, P.-T. *et al.* Bacterial colony from two-dimensional division to three-dimensional development. *PLoS One* **7**, e48098 (2012).
8. Duvernoy, M.-C. *et al.* Asymmetric adhesion of rod-shaped bacteria controls microcolony morphogenesis. *Nat. Commun.* **9**, 1120 (2018).
9. Cates, M. E. & Tjhung, E. Theories of binary fluid mixtures: from phase-separation kinetics to active emulsions. *J. Fluid Mech.* **836**, P1 (2018).
10. Beris, Antony. N. & Edwards, B. J. *Thermodynamics of flowing systems: with internal microstructure*. (Oxford University Press, 1994).
11. Doostmohammadi, A., Thampi, S. P. & Yeomans, J. M. Defect-mediated morphologies in growing cell colonies. *Phys. Rev. Lett.* **117**, 048102 (2016).

12. Dell'Arciprete, D. *et al.* A growing bacterial colony in two dimensions as an active nematic. *Nat. Commun.* **9**, 4190 (2018).
13. You, Z., Pearce, D. J. G., Sengupta, A. & Giomi, L. Geometry and mechanics of microdomains in growing bacterial colonies. *Phys. Rev. X* **8**, 031065 (2018).
14. Marchetti, M. C. *et al.* Hydrodynamics of soft active matter. *Rev. Mod. Phys.* **85**, 1143–1189 (2013).
15. Copenhagen, K., Alert, R., Wingreen, N. S. & Shaevitz, J. W. Topological defects promote layer formation in *Myxococcus xanthus* colonies. *Nat. Phys.* **17**, 211–215 (2021).
16. Bär, M., Großmann, R., Heidenreich, S. & Peruani, F. Self-propelled rods: insights and perspectives for active matter. *Annu. Rev. Condens. Matter Phys.* **11**, 441–466 (2020).
17. Meacock, O. J., Doostmohammadi, A., Foster, K. R., Yeomans, J. M. & Durham, W. M. Bacteria solve the problem of crowding by moving slowly. *Nat. Phys.* **17**, 205–210 (2021).
18. Volfson, D., Cookson, S., Hasty, J. & Tsimring, L. S. Biomechanical ordering of dense cell populations. *Proc. Natl. Acad. Sci. USA* **105**, 15346–15351 (2008).
19. You, Z., Pearce, D. J. G. & Giomi, L. Confinement-induced self-organization in growing bacterial colonies. *Sci. Adv.* **7**, eabc8685 (2021).
20. Başaran, M., Yaman, Y. I., Yuce, T. C., Vetter, R. & Kocabas, A. Large-scale orientational order in bacterial colonies during inward growth. (2021).
21. Cho, H. *et al.* Self-organization in high-density bacterial colonies: efficient crowd control. *PLoS Biol.* **5**, e302 (2007).
22. Sheats, J., Sclavi, B., Cosentino Lagomarsino, M., Cicuta, P. & Dorfman, K. D. Role of growth rate on the orientational alignment of *Escherichia coli* in a slit. *R. Soc. Open Sci.* **4**, (2017).

23. Thijssen, K., Metselaar, L., Yeomans, J. M. & Doostmohammadi, A. Active nematics with anisotropic friction: the decisive role of the flow aligning parameter. *Soft Matter* **16**, 2065–2074 (2020).

24. Liu, S., Shankar, S., Marchetti, M. C. & Wu, Y. Viscoelastic control of spatiotemporal order in bacterial active matter. *Nature* **590**, 80–84 (2021).

25. Drescher, K. *et al.* Architectural transitions in *Vibrio cholerae* biofilms at single-cell resolution. *Proc. Natl. Acad. Sci. USA* **113**, E2066-2072 (2016).

26. Yan, J., Sharo, A. G., Stone, H. A., Wingreen, N. S. & Bassler, B. L. *Vibrio cholerae* biofilm growth program and architecture revealed by single-cell live imaging. *Proc. Natl. Acad. Sci. USA* **113**, E5337–E5343 (2016).

27. Berk, V. *et al.* Molecular architecture and assembly principles of *Vibrio cholerae* biofilms. *Science* **337**, 236–239 (2012).
